# Calmodulin controls spatial and temporal specificity of calcium-induced calcium release

**DOI:** 10.64898/2026.02.11.705296

**Authors:** Joanna Jędrzejewska-Szmek, Kim T Blackwell

## Abstract

Calcium dynamics controls learning and memory, and abnormal calcium dynamics have been implicated in neurodegenerative disorders, such as Alzheimer’s disease (AD). Calcium dynamics are influenced by calcium-induced calcium release (CICR), which is mediated by ryanodine receptors (RyR) located on endoplasmic reticulum (ER) membrane. Calmodulin, one of the most abundant proteins in the brain, inhibits RyR2, expressed in the dendrites of hippocampal CA1 neurons, with several reported consequences: relief of this inhibition is responsible for heart failure, and enhancing calmodulin to RyR binding [1] alleviates cell loss and AD-like neuronal hyperexcitability. To investigate the role of calmodulin in aging and AD, we built a sophisticated reaction-diffusion model of a dendritic branch with ER. We showed that relieving calmodulin inhibition of RyR2 increased spatial and temporal spread of calcium transients in the dendrite. This effect was also visible in a model of old age, where disinhibition of half of the RyR2 population increased spatial spread of calcium transients by a factor of 2, and disinhibition of RyR2 combined with increased concentration of calcium buffering molecules increased duration of calcium transients. Lower activation of plasma membrane calcium ATPase (PMCA), which is also activated by calmodulin and inhibited by *β*-Amyloid oligomers, and not RyR2 disinhibition, led to an increase in resting intracellular calcium concentration as observed in AD. Overall, our research demonstrates that changes in calmodulin that are associated with AD and aging, by regulation of RyR2 (in old age) and PMCA (in AD), underlie changes in calcium dynamics that might have consequences for learning and memory.

**Author summary:** Calcium dynamics controls learning and memory. In neurons calcium dynamics are regulated by a complex system including calcium-permeable channels and extrusion pumps. The components of this system are regulated by calmodulin, which is enriched in the brain. We show that relieving calmodulin inhibition of calcium permeable channels, which accompanies aging, decreases spatial and temporal specificity of calcium release, contributing to the deficits in learning and memory observed in old age. In contrast, calcium extrusion pumps that are activated by calmodulin are likely responsible for increased resting intracellular calcium concentration observed in Alzheimer’s disease. In consequence, a novel role emerges for calmodulin, which in the brain is primarily considered a fast-acting calcium buffering molecule, as an important regulator in neuronal calcium dynamics underlying pathology.

## Introduction

In many neurodegenerative disorders such as Alzheimer’s disease (AD), neural information processing is deficient, interfering with a patient’s daily life [2]. Neuronal information processing begins in the dendrites [3], which continuously receive information from other neurons in the form of synaptic activation. Dendritic calcium dynamics, one of the modes of information processing [4] in the nervous system, has been extensively studied, with a focus on activation of N-methyl-D-aspartate (NMDA) receptors [5] and voltage-gated calcium permeable channels (VGCC) [3]. However, relatively little is known about the role of calcium stores such as the endoplasmic reticulum (ER) in information processing in the dendrite [6]. It is crucial to investigate the role of the ER in calcium dynamics since its calcium-release channels have been implicated in development and alleviation of Alzheimer’s disease in murine models [1, 7, 8].

The ER is a large intracellular organelle with continuous membranes and can be viewed as a cable-within-the neuron [9]. Calcium is released from the ER through inositol triphosphate (IP_3_) receptors (IP_3_R) in response to IP_3_ and ryanodine receptors (RyR) in response to calcium elevation [6]. The latter process is known as calcium-induced calcium release (CICR) and serves as an amplifier of cytosolic calcium signals.

There are 3 RyR isoforms, which are expressed in the brain [10], with RyR2 expressed in the dendritic shafts [11] and RyR3 expressed in the spines [12] of hippocampal CA1 pyramidal neurons. RyRs are ion channels that are modulated by many proteins such as calmodulin (CaM), calcium/calmodulin activated protein kinase II and protein kinase A. Calmodulin in the brain is mostly viewed as a fast acting calcium buffering molecule [13]. However, it is a vital part of the calcium signal transduction pathway, not only activating the calcium/calmodulin activated protein kinases, but also activating RyR1 and RyR3 isoforms and inhibiting RyR2. Moreover, due to its abundance, calmodulin might be constitutively bound to RyRs in the brain [14, 15]. Yet, with oxidative stress and aging the affinity of calmodulin to RyR is lowered [16], possibly relieving the inhibition of RyR2 by calmodulin. Disinhibited RyR2s might contribute to aberrant calcium homeostasis [6] that is responsible for disruption of neuronal function in AD [7, 17] and deficits in learning and memory accompanying old age. Indeed, enhancing calmodulin binding to RyR2 in mouse models of AD reversed cognitive dysfunction [1].

The role of CICR in information transfer along the dendrite is unclear. Information transfer from the synapse to the nucleus and back might involve electrotonic propagation involving the ER along the dendritic tree [9], but experiments performed in cultured hippocampal neurons show that ER membrane potentials do not spread beyond the site of activation [18]. Another possibility of biochemical signaling to the nucleus is induction of an intracellular calcium wave propagating down the dendritic tree. Excitation-translation coupling in neuronal cultures depends on activation of L-type VGCCs [19, 20] and also recruits RyR2s [19, 21]. CICR contributes to the propagation of dendritic transients [22] and RyR2 activation contributes to nuclear calcium release [21]. It has recently been suggested that RyR2s might support propagation of intracellular calcium waves evoked by back-propagating action potentials (bAP) in apical dendrites [23]. Yet, experimental evidence indicates that intracellular calcium waves in dendrites of pyramidal neurons in-vitro cannot be evoked by bAPs [24], and require activation of metabotropic glutamate receptors, which in turn leads to IP_3_ production and IP_3_-induced calcium release [25–27], which was investigated computationally in [28, 29].

To test whether RyR2s alone can support calcium wave propagation, as suggested by [23], we built spatial stochastic reaction-diffusion models of hippocampal CA1 pyramidal thin, medium and thick dendrites. We show in-silico that RyR2s **cannot** support intracellular calcium wave propagation in a model of a healthy neuron (control conditions) **nor** when calmodulin inhibition of RyR2s is relieved. Furthermore, we show that disinhibition of RyR2 increases spatial spread of calcium transients (lowers spatial specificity) and increases duration of calcium transients (decreases temporal specificity) in the dendrite. Using a model of CA1 pyramidal dendrites in old age we show that spatial and temporal specificity of calcium elevations is diminished in old age due to oxidization of calmodulin and RyR2s, possibly contributing to neuronal hyperexcitability observed in aging and AD [30, 31]. Finally, we show that calmodulin oxidation can be responsible for increased basal intracellular calcium concentration by lowering plasma membrane Ca^2+^ ATPase (PMCA) activity but not by RyR2-disinhibition.

## Materials and methods

### Computational methods

To investigate the extent of calcium elevation in the dendritic branch following synaptic stimulation we developed a multi-compartment stochastic reaction-diffusion model of signaling pathways underlying calcium-induced calcium release. We adapted the calcium regulation pathways implemented in [32], and added a model of the ER containing: luminal calcium (Ca_ER_) and calreticulin (CRT) with sarco/endoplasmic reticulum calcium ATPase (SERCA) and a leak channel (LeakER). The ER contained ryanodine receptors, as explained below, and store operated calcium entry (SOCE) [33], with parameters fitted to measurements from [34]. The ER was treated as a separate compartment that is uniformly distributed in the dendrite, occupying 25 % of its volume. Tortuosity of the ER was taken into account by reducing the diffusion rate of calcium molecules in the ER, compared to diffusion rate of the cytosolic calcium molecules. The signaling pathways model diagram is shown in Fig 1.

**Fig 1.**
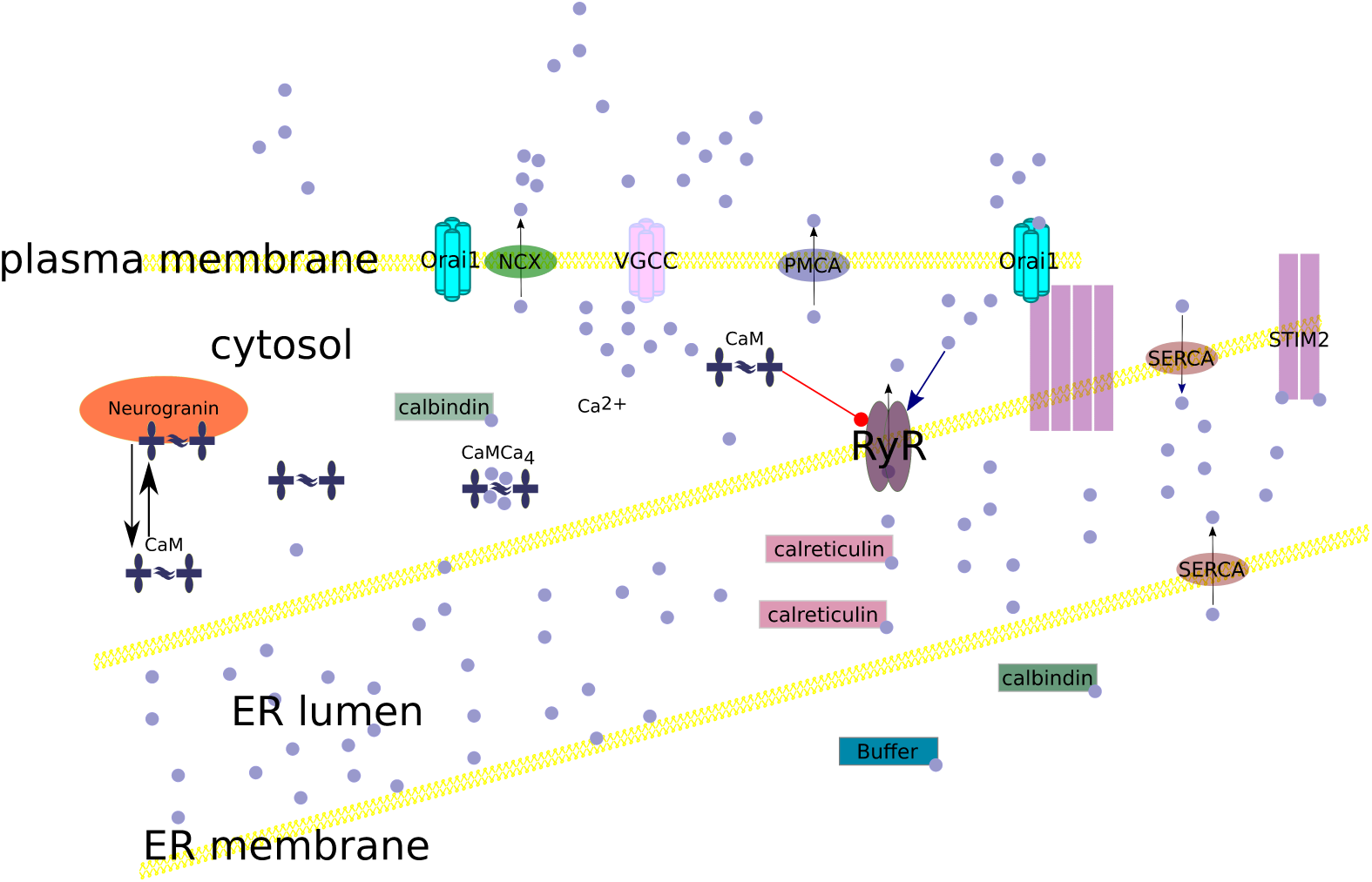
Molecular pathways implemented in the model. Calcium flows into the cell through the calcium permeable channels in the cell membrane (denoted VGCC in the diagram and Leak in Tab. 1). Calcium is extruded into the extracellular space by the plasma membrane Ca^2+^-ATPase (PMCA) and sodium calcium exchanger (NCX) and into the endoplasmic reticulum by the Sarco/endoplasmic Reticulum Calcium ATPase (SERCA). In the cytosol calcium is buffered by calbindin, indiffusible buffer (Buffer) and calmodulin (CaM). Apo-calmodulin (unbound to calcium) is buffered by neurogranin (Ng). Calcium flows from the ER to the cytosol through open ryanodine type 2 receptors (RyR2), which are activated by calcium and inhibited by calmodulin. In the ER calcium is buffered by calreticulin (CRT) and binds stromal interaction molecules (STIM), which serve as luminal calcium sensors. When calcium in the endoplasmic reticulum is too low, STIM molecules form dimers and bind calcium release-activated calcium channel protein 1 (Orai1) opening them and allowing calcium from the extracellular space to flow into the cytosol, where it would be transferred by SERCA to the endoplasmic reticulum, thereby replenishing it. The ORA1-STIM channels constitute store-operated calcium entry (SOCE).

**Table 1.** Calcium extrusion to the extracellular space and calcium buffering in the cytosol: reaction rates.

| Reaction | $k_f$ ($\text{nM}^{-1}\text{ms}^{-1}$) | $k_r$ ($\text{ms}^{-1}$) | $k_{\text{cat}}$ ($\text{ms}^{-1}$) | source |
| --- | --- | --- | --- | --- |
| $\text{PMCA} + \text{Ca}^{2+} \rightleftharpoons \text{PMCA} \cdot \text{Ca} \rightarrow \text{PMCA} + \text{Ca}_{\text{ext}}^{2+}$ | 0.5e-4 | 0.007 | 0.0035 | [40] |
| $\text{NCX} + \text{Ca}^{2+} \rightleftharpoons \text{NCX} \cdot \text{Ca} \rightarrow \text{NCX} + \text{Ca}_{\text{ext}}^{2+}$ | 1.68e-5 | 0.0112 | 0.0056 | [41, 42] |
| $\text{Leak} + \text{Ca}_{\text{ext}}^{2+} \rightleftharpoons \text{Leak} \cdot \text{Ca}_{\text{ext}} \rightarrow \text{Leak} + \text{Ca}^{2+}$ | 1.5e-6 | 1.1e-3 | 1.1e-3 | adjusted to maintain $\text{Ca}_i^{2+}$ concentration of 75 nM [32], represents VGCCs |
| $\text{Calbindin} + \text{Ca}^{2+} \rightleftharpoons \text{CalbindinCa}$ | 2.8e-5 | 0.0196 | | [43] |
| $\text{IndiffusibleBuffer} + \text{Ca}^{2+} \rightleftharpoons \text{IndiffusibleBufferCa}$ | 0.004e-3 | 20.0e-2 | | [44] |
| $2\text{Ca}^{2+} + \text{CaM} \rightleftharpoons \text{CaM}\text{Ca}_2$ | 6e-6 | 9.1e-3 | | [45] |
| $2\text{Ca}^{2+} + \text{CaM} \rightleftharpoons \text{CaM}\text{Ca}_2$ | 0.1e-3 | 1000e-3 | | [46] |
| $\text{CaM} + \text{Ng} \rightleftharpoons \text{NgCaM}$ | 2.80E-5 | 3.60E-2 | | [47] |

In the model ER molecules were distributed randomly according to a mean concentration. SERCA and STIM_2_ concentrations were scaled by area/volume ratio of the dendrite. RyR2 copy numbers were kept the same for all dendritic diameter models, because experimental evidence suggests that their concentration depends linearly on the total length of the dendrite [35]. ER luminal species (Ca_ER_, CRT and Ca-bound calreticulin (CRT · Ca_ER_)) were proportional to ER volume with concentration set to achieve ER molecule copy numbers expected from published measurements of concentration [36].

In our simulations we used two models of ryanodine receptors, which were based on the Keizer-Smith model [37]:

1. RyR2CaM – a model of ryanodine receptor constitutively bound to calmodulin (inhibited by calmodulin) and activated by calcium, fitted to data presented in Fig 2 of [38],
2. RyR2 no CaM – a model of ryanodine receptor not bound to calmodulin (disinhibited) as occurs when calmodulin is oxidized, which was fitted to data presented in Fig 2 of [39].

**Fig 2.**
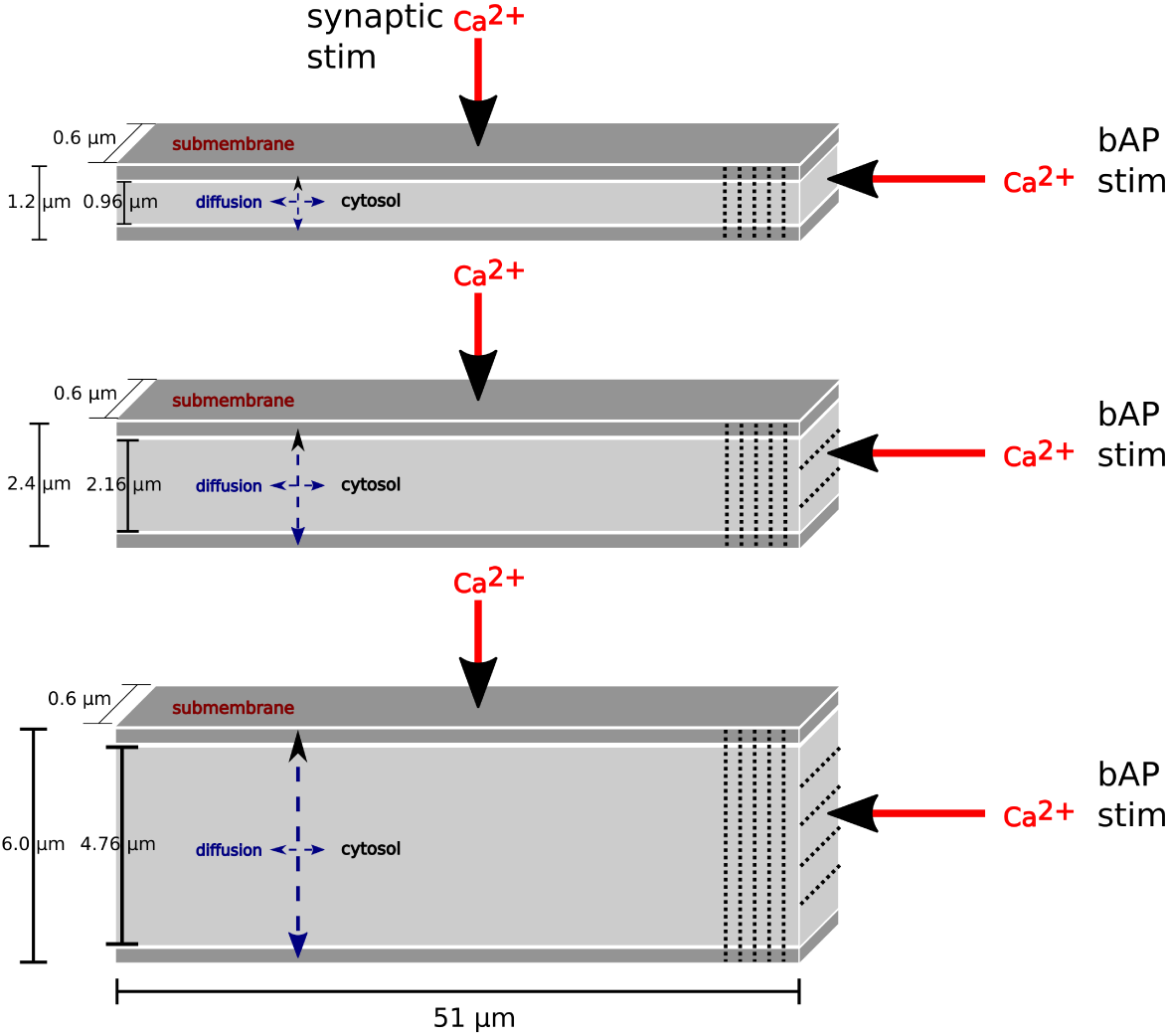
Dendritic morphologies used in simulations. We used three dendritic morphologies: thin dendrite (diameter of 1.2 µm), medium dendrite (diameter of 2.4 µm) and thick dendrite (diameter of 6.0 µm). All dendrites were 51 µm long. Each morphology was subdivided into voxels, allowing two-dimensional diffusion. Submembrane voxels were cubes of 0.5 µm × 0.4 µm × 0.6 µm (the same for all dendritic diameters). The submembrane voxels allow for placement of membrane associated molecules. Cytosolic voxels were the same as submembrane voxels for the 1.2 µm model, cubes of 0.5 µm × 0.46 µm × 0.6 µm for the 2.4 µm model and cubes of 0.5 µm × 0.4615 µm × 0.6 µm for the 6.0 µm model. The dotted lines show some of the lengthwise and widthwise subdivisions into voxels. For simulating synaptic activation, calcium molecules were injected into submembrane voxels in the middle of the dendritic morphology (denoted synaptic stim on the diagram). For simulating b-AP like stimulus, calcium molecules were injected into the cytosol to an end dendritic voxel (point stimulus, denoted bAP stim on the diagram).

### Morphology

We implemented the signaling pathways model (Fig 1) in 51 µm long dendrites with diameters of 1.2 µm, 2.4 µm and 6.0 µm representing thin, medium and thick dendrites, respectively (Fig 2). Each morphology was divided into voxels. Submembrane voxels, which allow for placing of membrane-associated molecules, were cubes of 0.5 µm × 0.4 µm × 0.6 µm (the same for all dendritic diameters). Cytosolic voxels were the same as submembrane voxels for the 1.2 µm model, cubes of 0.5 µm × 0.46 µm × 0.6 µm for the 2.4 µm model and cubes of 0.5 µm × 0.4615 µm × 0.6 µm for the 6.0 µm model.

### Control model

The control model comprised a model of cytosolic calcium dynamics (Tab 1) with total concentrations shown in Tab 2 and diffusible species in Tab 4. This model was fitted with the model of the ER containing calcium extrusion and buffering (Tab 3), opening and calcium release by the RyR2 inhibited by calmodulin (upper half of Tab 5) and SOCE (Tab 6) with total concentrations specified in Tab 7.

### Old age model

Aging is related to changes in calcium homeostasis [63]. Calmodulin becomes oxidized [64], which reduces its binding to RyR2 [16]. Specifically, in the oxidized conditions with cytosolic calcium levels calmodulin affinity for RyR2 drops twenty-fold [65], resulting in 50% RyR2s bound to calmodulin and hence inhibited by calmodulin. PMCA activity is diminished [66–68] and in in-vitro aging cultures, store-operated calcium entry is down-regulated [69]. The buffering capacity of intracellular buffers of CA1 pyramidal neurons is increased [70]. Also the RyR2 expression is increased [71]. The T-type calcium channels are down-regulated in the hippocampus of aging mice and men [72], whereas the L-type channels are up-regulated [73]. It has also been reported that the resting intracellular calcium concentration in CA1 pyramidal models from aged rats is unchanged [48]. To take all these changes into account in the old age model we modified the control model in several ways.

1. We lowered PMCA *k*_cat_ by 20% (Table 8).
2. To emulate changes in calmodulin binding to RyR2 due to oxidation, we divided the population into halves: one bound to and, in consequence, inhibited by calmodulin (upper half of Tab 5) and one not inhibited by calmodulin (lower half of Tab 5), as indicated by Table 7. This resulted in the total RyR2 population favoring activation by calcium consistent with experimental data [74].
3. To account for increased calcium buffering in old age we doubled calbindin and indiffusible buffer concentrations (Table 2).
4. To account for an upregulation of L-type VGCCs we increased calcium inputs (a comparison of calcium inputs is shown Tab 9).

**Table 2.** Concentrations of molecular species governing cytosolic calcium dynamics (Table 1): calcium extrusion (PMCA and NCX) and calcium buffering (calmodulin, calbindin, indiffusible calcium buffer) molecules. Concentration units of cytosolic species ((Ca^2+^), Ca_ext_^2+^, calbindin, calmodulin, neurogranin, indiffusible calcium buffer, ER calcium and calreticulin) are nM, whereas densities of membrane species (Leak, PMCA, NCX) are expressed in pmol µm^−2^.

| Specie | control | disinhibited<br>RyR2 | old age | AD | source |
| --- | --- | --- | --- | --- | --- |
| calcium<br>($\text{Ca}^{2+}$) | 75 | 75 | 75 | | [48, 49] |
| extracellular<br>calcium<br>($\text{Ca}_{\text{ext}}^{2+}$) | $2 \times 10^6$ | $2 \times 10^6$ | $2 \times 10^6$ | $2 \times 10^6$ | chosen to match in-vitro experiments |
| Leak | 6000 | 6000 | 5000 | 6000 | chosen to counterbalance PMCA |
| PMCA | 5600 | 5600 | 5600 | 5600 | [50] |
| NCX | 3000 | 3000 | 3000 | 3000 | adjusted for $V_{\text{max}}$ of NCX to match [51] |
| Calbindin | $150 \times 10^3$ | $150 \times 10^3$ | $300 \times 10^3$ | $150 \times 10^3$ | to match 160 $\mu\text{M}$ high affinity binding sites [52] |
| Calmodulin | $25.8 \times 10^3$ | $25.8 \times 10^3$ | $23.0 \times 10^3$ | $25.8 \times 10^3$ | [14] |
| Neurogranin<br>(Ng) | $20 \times 10^3$ | $20 \times 10^3$ | $40 \times 10^3$ | $20 \times 10^3$ | [53] |
| Indiffusible<br>calcium<br>buffer | $2 \times 10^6$ | $2 \times 10^6$ | $4 \times 10^6$ | $2 \times 10^6$ | [44] |
| ER cal-<br>cium<br>($\text{Ca}_{\text{ER}}^{2+}$) | $320 \times 10^3$ | 320 | 320 | 320 | [54] |
| calreticulin<br>(CRT) | $400 \times 10^3$ | $400 \times 10^3$ | $400 \times 10^3$ | $400 \times 10^3$ | [36] |

**Table 3.**
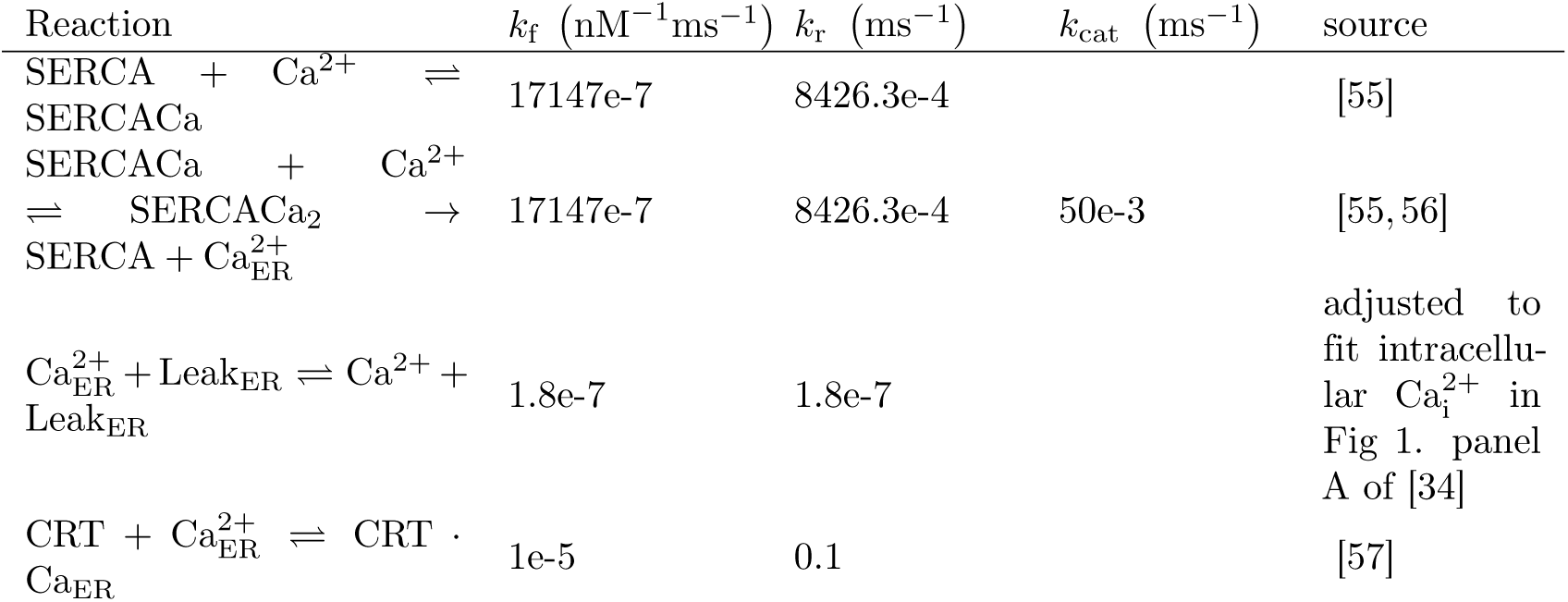
Calcium extrusion to the ER and calcium buffering in the ER: reaction rates.

**Table 4.** Diffusion rates of diffusible species. Abbreviations same as in Table 1, Table 3,Table 6, Table 5.

| Name | $k_{\text{diff}}$ ($\mu\text{m}^2 \text{s}^{-1}$) | source |
| --- | --- | --- |
| $\text{Ca}^{2+}$ | 100 | [58] |
| ext. $\text{Ca}^{2+}$ | 10 | [32] |
| ER $\text{Ca}^{2+}$ | 10 | Lower than cytosolic diffusion to account for complex structure of ER tubules |
| CaM | 4 | [59] |
| Calbindin | 8.3 | [60] |

**Table 5.** Model of calmodulin-bound RyR2 (upper half) and disinhibited RyR2 (lower half). Both models were based on [37], with calcium binding constants fitted to data in Fig 2 [38] (+1 µM CaM) for the RyR2CaM model, and Fig 2 of [39] for the RyR2 model.

| Reaction | $k_f$ ($\text{nM}^{-4} \text{ms}^{-1}$) | $k_r$ ($\text{ms}^{-1}$) | source |
| --- | --- | --- | --- |
| $\text{RyR2CaM} + 4\text{Ca}^{2+} \rightleftharpoons \text{RyR2CaMC}_1$ | 10e-11 | 1 | [38] |
| $\text{RyR2CaMC}_1 + 4\text{Ca}^{2+} \rightleftharpoons \text{RyR2CaMO}_1$ | 5e-10 | 4.8 | [38] |
| $\text{RyR2CaMO}_1 + 4\text{Ca}^{2+} \rightleftharpoons \text{RyR2CaMO}_2$ | 5e-12 | 13 | [38] |
| $\text{RyR2CaMC}_2 + 4\text{Ca}^{2+} \rightleftharpoons \text{RyR2CaMO}_2$ | 3.33333e-16 | 66.667e-2 | [38] |
| $\text{RyR2CaMC}_1 + 4\text{Ca}^{2+} \rightleftharpoons \text{RyR2CaMC}_2$ | 1e-9 | 1.248e-2 | [38] |
| $\text{RyR2CaMC}_2 + 4\text{Ca}^{2+} \rightleftharpoons \text{RyR2CaMC}_3$ | 1.5e-15 | 3 | [38] |
| $\text{RyR2CaMO}_1 + \text{Ca}_{\text{ER}}^{2+} \rightleftharpoons \text{RyR2CaMO}_1 + \text{Ca}^{2+}$ | 5e-3 | 5e-3 | [61] |
| $\text{RyR2CaMO}_2 + \text{Ca}_{\text{ER}}^{2+} \rightleftharpoons \text{RyR2CaMO}_2 + \text{Ca}^{2+}$ | 5e-3 | 5e-3 | [61] |
| $\text{RyR2} + 4\text{Ca}^{2+} \rightleftharpoons \text{RyR2C}_1$ | 5e-11 | 5e-3 | [39] |
| $\text{RyR2C}_1 + 4\text{Ca}^{2+} \rightleftharpoons \text{RyR2O}_1$ | 5e-10 | 9.6 | [39] |
| $\text{RyR2O}_1 + 4\text{Ca}^{2+} \rightleftharpoons \text{RyR2O}_2$ | 5e-11 | 13 | [39] |
| $\text{RyR2C}_2 + 4\text{Ca}^{2+} \rightleftharpoons \text{RyR2O}_2$ | 3.33333e-15 | 66.667e-3 | [39] |
| $\text{RyR2C}_1 + 4\text{Ca}^{2+} \rightleftharpoons \text{RyR2C}_2$ | 5000e-15 | 1.235e-3 | [39] |
| $\text{RyR2C}_2 + 4\text{Ca}^{2+} \rightleftharpoons \text{RyR2C}_3$ | 1e-14 | 3 | [39] |
| $\text{RyR2O}_1 + \text{Ca}_{\text{ER}}^{2+} \rightleftharpoons \text{RyR2O}_1 + \text{Ca}^{2+}$ | 5e-3 | 5e-3 | [61] |
| $\text{RyR2O}_2 + \text{Ca}_{\text{ER}}^{2+} \rightleftharpoons \text{RyR2O}_2 + \text{Ca}^{2+}$ | 5e-3 | 5e-3 | [61] |

**Table 6.** Store operated calcium entry model: reaction rates. ER calcium to STIM dimers binding rates were adjusted to fit intracellular Ca_i_^2+^ in Fig 1. panel A of [34]. Calcium flux through the open Orai1 channels was adjusted to fit intracellular Ca_i_^2+^ in Fig 1. panel A of [34] and the graded opening of SOCE by activation of different numbers of Orai1 subunits [62].

| Reaction | $k_f$ ($\text{nM}^{-1}\text{ms}^{-1}$) | $k_r$ ($\text{ms}^{-1}$) | source |
| --- | --- | --- | --- |
| $\text{STIM}_2 + \text{Ca}_{\text{ER}}^{2+} \rightleftharpoons \text{STIM}_2\text{Ca}_{\text{ER}}$ | 8e-7 | 2e-3 | adjusted to fit intracellular $\text{Ca}_i^{2+}$ in Fig 1. panel A of [34] |
| $\text{STIM}_2 + \text{STIM}_2 \rightleftharpoons \text{STIM}_2 \text{ dimer}$ | 4.8e-7 | 11e-5 | [33] |
| $\text{PMJ} + \text{STIM}_4 \rightleftharpoons \text{PMJ} + \text{PMJSTIM}_4$ | 1.8e-04 | 3e-4 | [33] |
| $\text{PMJSTIM}_4 + \text{Orai1} \rightleftharpoons \text{Orai1STIM}_4$ | 1.5e-04 | 8e-3 | [33] |
| $\text{Orai1STIM}_4 + \text{Orai1STIM}_4 \rightleftharpoons \text{Orai1}_2\text{STIM}_4$ | 3.75e-5 | 3.75e-5 | [33,34] |
| $\text{Orai2STIM}_4 + \text{Orai1STIM}_4 \rightleftharpoons \text{Orai3STIM}_4$ | 9.375e-06 | 5e-4 | [33,34] |
| $\text{Orai3STIM}_4 + \text{Orai1STIM}_4 \rightleftharpoons \text{2Orai2STIM}_4$ | 9.375e-06 | | [33,34] |
| $\text{Ca}_{\text{ext}} + \text{Orai1STIM}_4 \rightleftharpoons \text{Orai1STIM}_4 + \text{Ca}^{2+}$ | 1.5e-11 | 1.5e-11 | [34,62] |
| $\text{Ca}_{\text{ext}} + \text{Orai2STIM}_4 \rightleftharpoons \text{Orai2STIM}_4 + \text{Ca}^{2+}$ | 5e-8 | 5e-8 | [34,62] |
| $\text{Ca}_{\text{ext}} + \text{Orai3STIM}_4 \rightleftharpoons \text{Orai3STIM}_4 + \text{Ca}^{2+}$ | 5e-8 | 5e-8 | [34,62] |

**Table 7.**
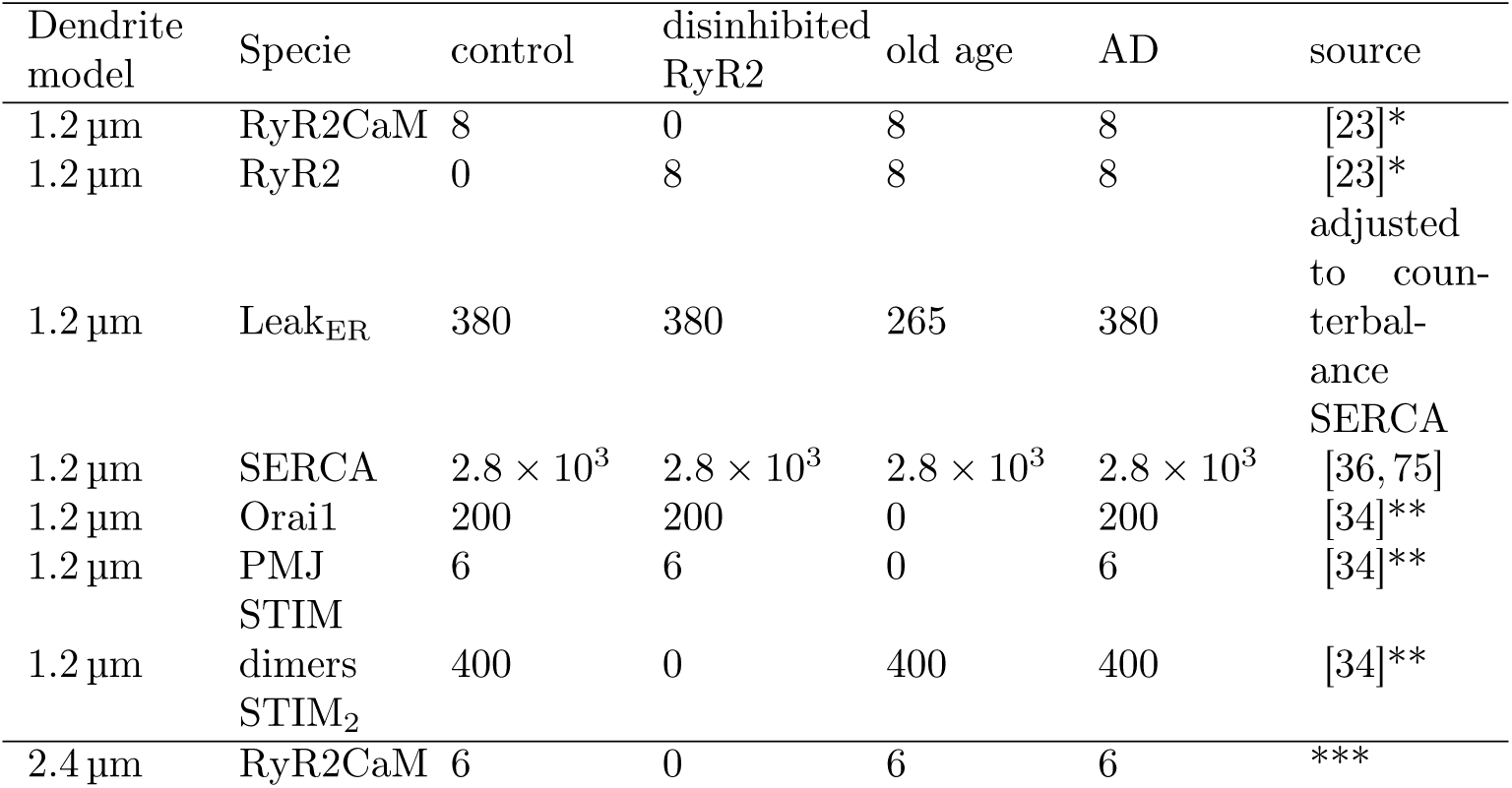
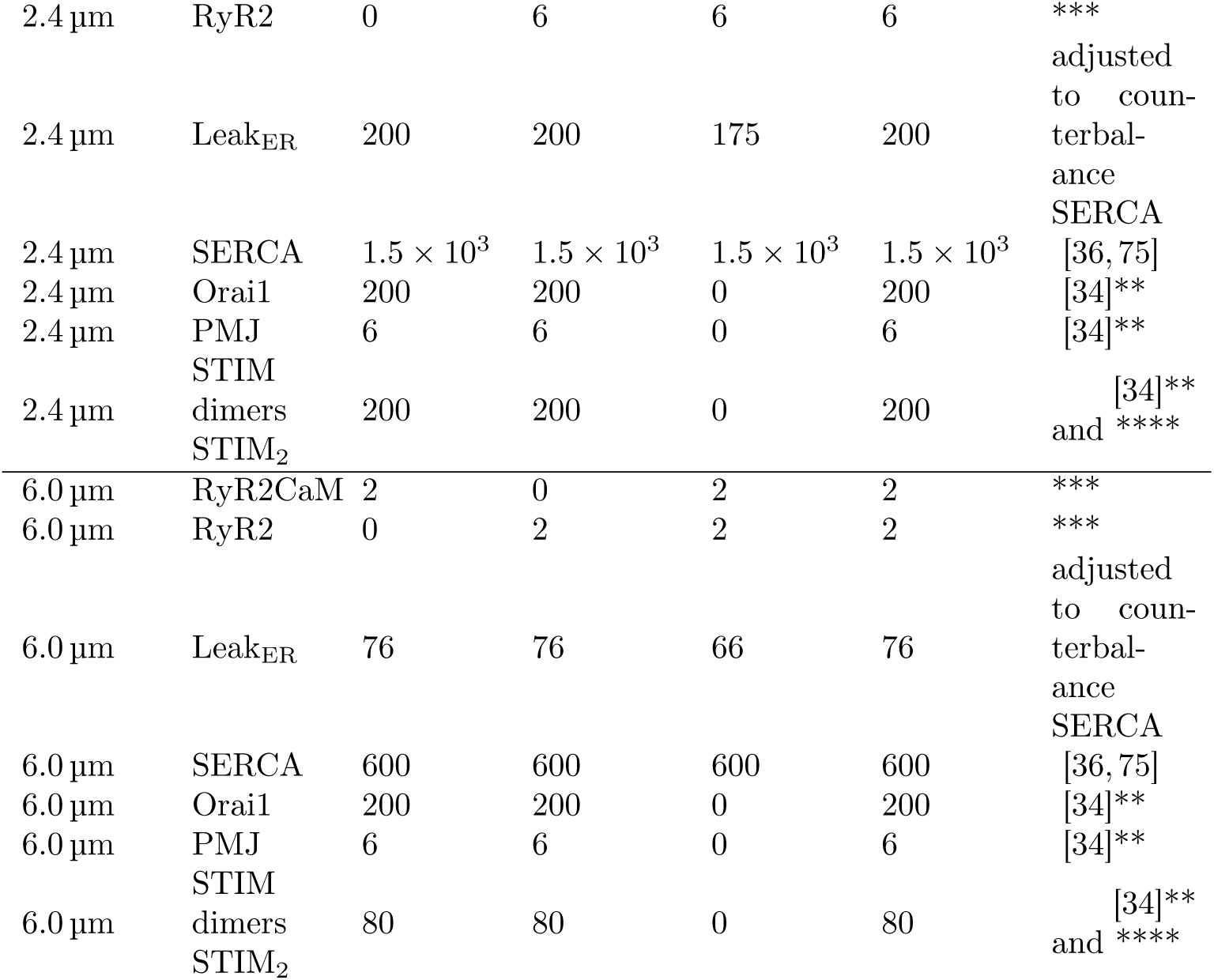
Total concentration of molecular species localized in the ER membrane. We assumed that the ER comprises 25% of the cytosol with ryanodine receptors (RyR2 and RyR2CaM), SERCA, Leak_ER_, and STIM dimers uniformly distributed in the cytosol with concentration units of nM, and Orai1 and plasma membrane-endoplasmic reticulum junctions (PMJ) located in dendritic membrane with units of pmol µm^−2^. *adjusted to match RyR2 density from [23]. **adjusted to fit intracellular Ca_i_^2+^ in Fig 1. panel A of [34]. ***adjusted to match RyR2 copy numbers for the 1.2 model. ****scaled by area-to-volume ratio of the ER.

### AD model

Alzheimer’s disease is characterized by calcium dishomeostasis [76] and elevated resting intracellular calcium concentration [77, 78]. Animal models of AD exhibit increased RyR expression [79, 80], high levels of oxidative stress [81], and reduced calbindin levels compared to old age [82]. High levels of oxidative stress likely result in high calmodulin oxidation resulting in lower PMCA activity and lower RyR2 inhibition. To investigate how PMCA and RyR2 influence resting intracellular calcium dynamics in early onset AD, we built an AD model of the dendrite (Tab 2 and Tab 7). This model exhibited lower PMCA activity (Tab 8), increased RyR2 concentration (Tab 7) with half the RyR2 population inhibited by calmodulin (upper half of Tab 5) and half disinhibited (lower half of Tab 5). In contrast to the old-age model, it did not have altered calcium buffering and it preserved calcium leak values of the control model.

**Table 8.**
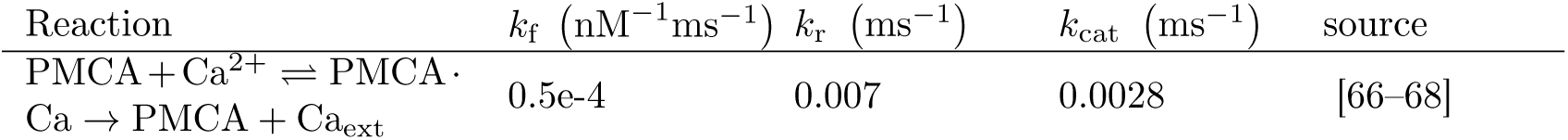
Calcium extrusion to the extracellular space reaction rates in the old age model.

### Stimulation protocols

All model variants were simulated without any stimulation to reach steady state (usually for 1000 s). Initial conditions of all the model species were recalculated based on the last 25 s of those simulations and the models were rerun without any stimulation to verify the stability of the steady state. If the steady state was not reached the procedure was repeated. For simulations of calcium wave induction only models in steady state were used.

We simulated two types of calcium influx to the dendrite: (Fig 2)

1. synaptic-like activation as a 40 ms influx of calcium molecules into submembrane voxels in the middle of the dendrite (summary of the stimulation is presented in Tab. 9),
2. bAP-like stimulus as a 3 ms influx of calcium molecules into the cytosol voxels at the end of the dendrite similarly as in [23] (summary of the stimulation is presented in Tab. 10).

**Table 9.**
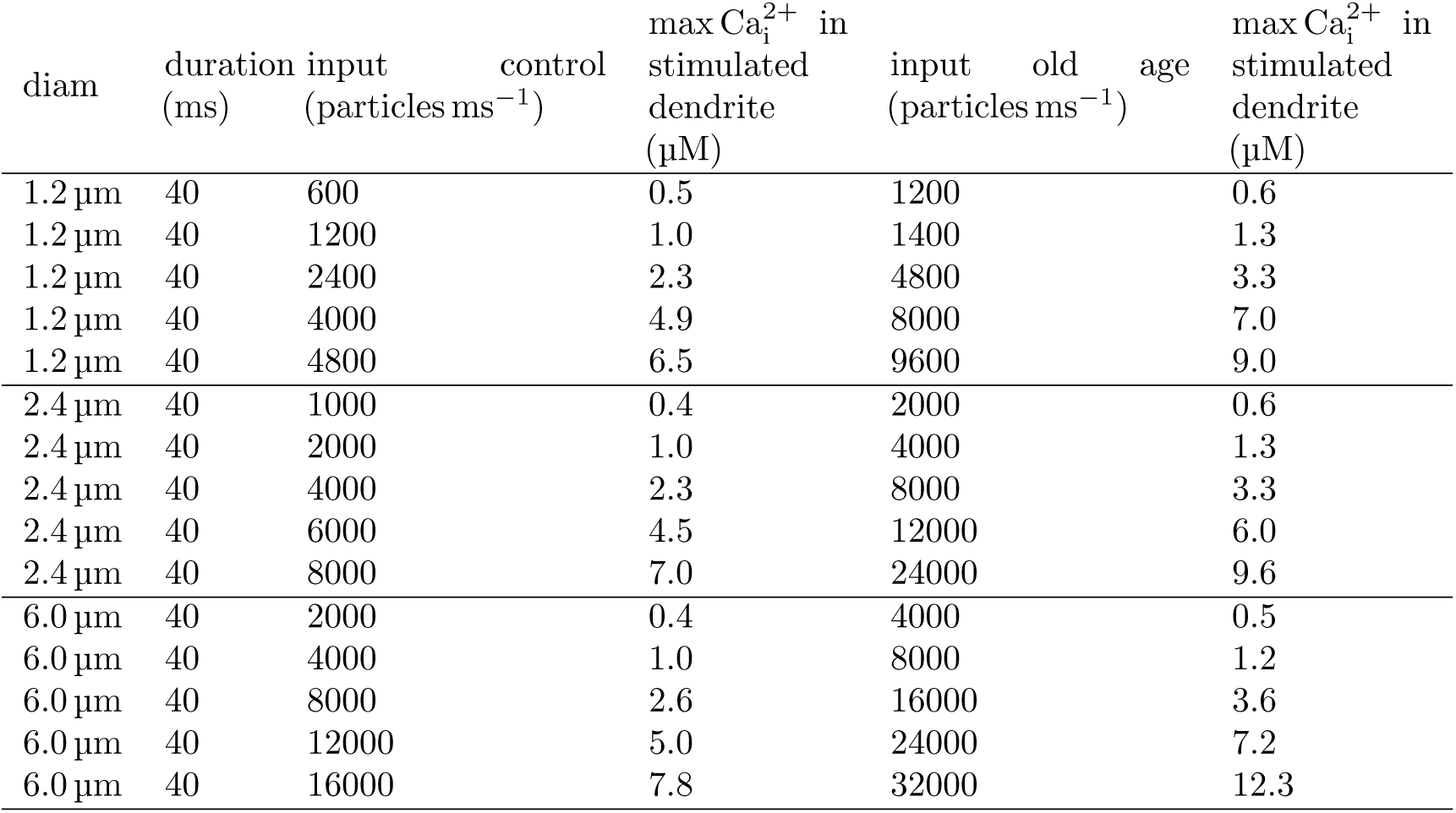
Calcium inputs for Fig 3, 5, 6. Stimulation was applied at 3000 ms to the submembrane voxels in the middle of the morphology.

**Table 10.** Calcium inputs for Fig 4. Stimulation was applied at 5000 ms. Stimulation was applied to the cytosol of the first dendritic segment.

| diam | duration (ms) | input (particles ms<sup>-1</sup>) | max Ca<sub>i</sub><sup>2+</sup> in stimulated dendrite (μM) |
| --- | --- | --- | --- |
| 1.2 μm | 3 | 6000 | 3.4 |
| 1.2 μm | 3 | 12000 | 7.7 |
| 1.2 μm | 3 | 24000 | 19.2 |
| 1.2 μm | 3 | 40000 | 41.0 |
| 1.2 μm | 3 | 48000 | 54.9 |
| 2.4 μm | 3 | 10000 | 3.4 |
| 2.4 μm | 3 | 20000 | 8.0 |
| 2.4 μm | 3 | 40000 | 21.1 |
| 2.4 μm | 3 | 60000 | 41.2 |
| 2.4 μm | 3 | 80000 | 69.6 |
| 6.0 μm | 3 | 20000 | 3.1 |
| 6.0 μm | 3 | 40000 | 8.2 |
| 6.0 μm | 3 | 80000 | 27.0 |
| 6.0 μm | 3 | 120000 | 56.7 |
| 6.0 μm | 3 | 160000 | 89.9 |

We used 5 different calcium input values to thoroughly explore CICR in the dendrite. It is estimated that calcium concentration in the dendritic spine due to an excitatory postsynaptic potential (EPSP) can reach up to 12 µM and 1 µM due to a bAP [83]. Calcium levels in the thin apical dendrites and dendritic spines are highly correlated [84], hence we chose calcium inputs resulting in an elevation of calcium concentration in the stimulated dendrite up to 10 µM (Tab. 9). For simulation of the bAP we selected injections of similar numbers of calcium molecules as with the EPSP though in a much smaller time window, to fully explore CICR properties (Tab. 10). Calcium injections (Tab. 9 and 10) were calibrated to obtain similar calcium elevations in the dendrite for the 2.4 µm and 6.0 µm models.

Models were run for 3 s before synaptic-like activation (Tab. 9) and for 5 s before the b-AP like stimulus (Tab. 10). In total models were run for 10 s. We established that 10 s simulation duration was sufficient to observe changes in calcium dynamics by running simulations for 20 s first. For each condition (calcium injection, model variant) we ran 10 simulations with different seeds.

Simulations of the impact of RyR2 variants and PMCA variants on resting intracellular calcium concentration were performed in the absence of any stimulation for 1000 s. For each model we ran 4 simulations with different seeds.

### Simulation

The model was implemented using the stochastic reaction–diffusion simulator NeuroRD [85] version 3.2.3, using the adaptive (asynchronous *τ*-leaping) numerical method with tolerance of 0.01. All data and code used for running experiments and plotting is available on a GitHub repository at https://github.com/asiaszmek/stochastic_ER.

### Analysis

We calculated the mean intracellular calcium concentration every 0.5 µm in the longitudinal direction (the voxel width in the longitudinal direction was 0.5 µm) by averaging across the width. Spatial spread was calculated as the furthest location from the stimulated site, in which calcium amplitude following stimulation exceeds 250% of resting calcium concentration of baseline (prior to the stimulation). For synaptic activation-like stimulation, in which the middle of the dendrite was stimulated, we average the spatial spread in the left and right halves of the dendrite. Temporal decay constant was calculated by fitting an exponential function *a* · exp(−*^t^/b*) + *c* to the 1 sec long average calcium concentration with initial time (and *a*) corresponding to the time (and value) of maximum calcium concentration in the stimulated site. Time decay constant value was the *b* parameter obtained in the fit.

Figures 3, 4, 5 and 6 show mean values of the spatial spread and temporal decay constant with standard error as errors bars obtained in 10 simulations with different seeds.

**Fig 3.**
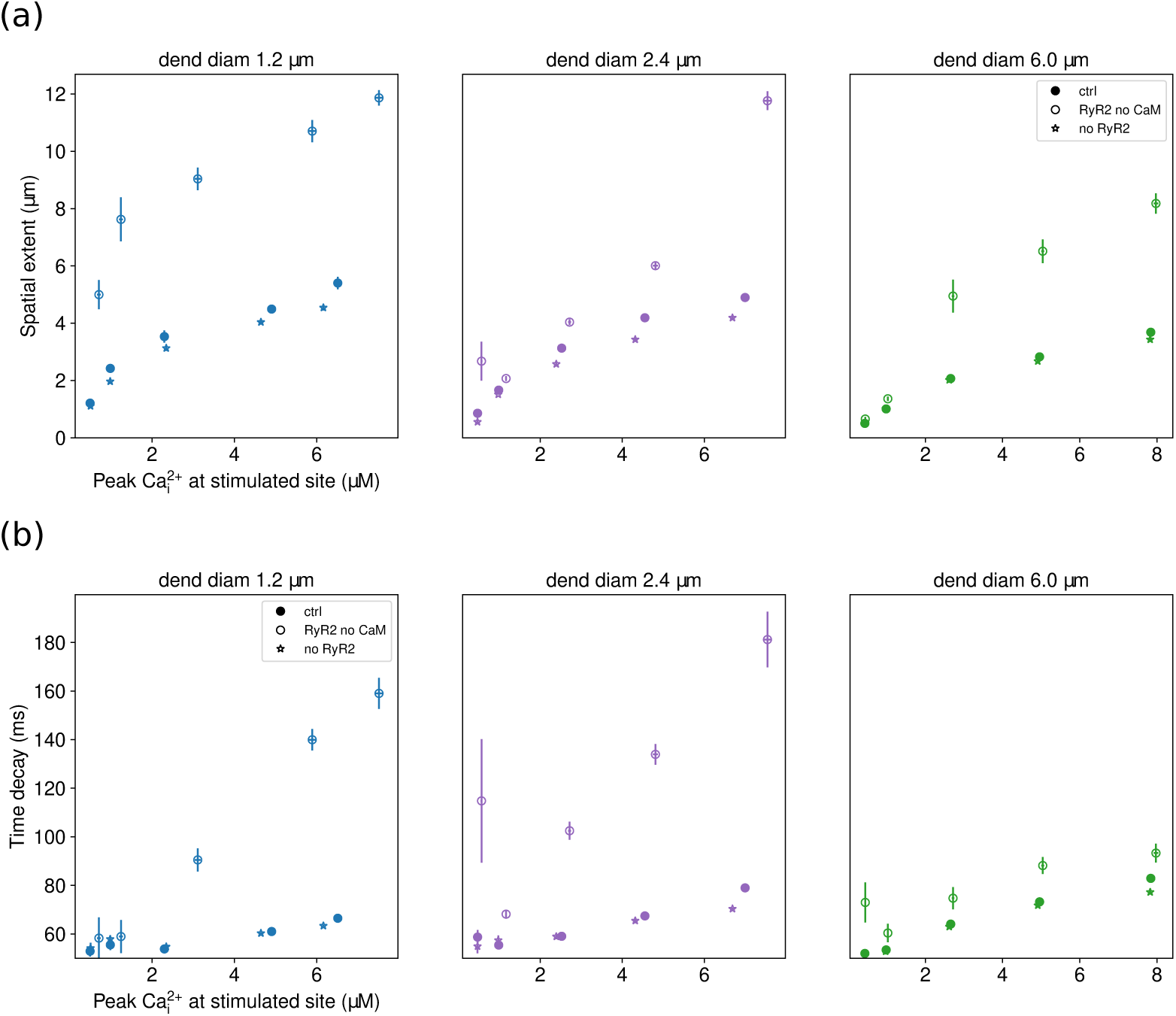
The ctrl model does not support intracellular calcium wave propagation in a CA1 pyramidal neuron dendrite model. Spatial extent (a) and temporal decay constant (b) of calcium transients for the model with RyR2s inhibited by CaM (ctrl), the model with disinhibited RyR2s (RyR2 no CaM) and the model without RyR2s (no RyR2). Disinhibition of RyR2 lowers spatial and temporal specificity of calcium transients (*p <* 0.001 for every panel of (a) and (b)). The horizontal error bars do not appear because they are smaller than the data points on the plots. With disinhibition of RyR2 (RyR2 no Cam) the spatial spread of calcium transients exceeded 5 µm. The spatial extent is the furthest location from stimulated site (middle of the dendrite) where calcium amplitude following stimulation exceeds 250% of resting calcium concentration. (b) CICR did not prolong calcium transients. Time decay constant was obtained by fitting a single exponential decay function to calcium concentration after the stimulation at the stimulated section of the dendrite.

**Fig 4.**
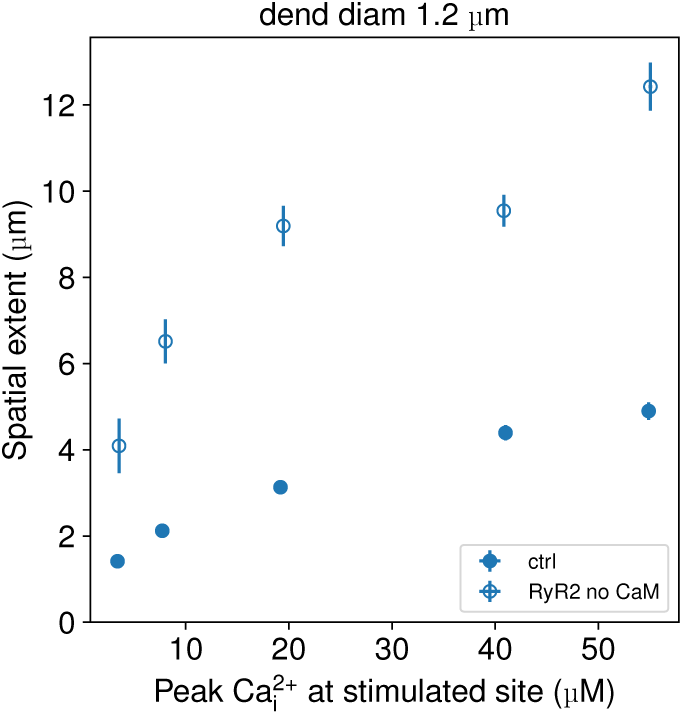
Spatial spread of calcium transients for a bAP-like stimulus. Only results for the thin dendrite are shown. A short calcium input (3 ms) was applied directly to the cytosol at one side of the dendrite as in [23] (also shown in Fig 2). Disinhibition of RyR2 by calmodulin (empty circles) increases spatial extent, but does not allow for calcium wave propagation. Spatial extent was the furthest location from the stimulated site where calcium amplitude following stimulation exceeded 250% of resting calcium concentration. The horizontal error bars do not appear because they are smaller than the data points on the plots.

**Table 11.** ANCOVA statistical significance results. Independent variables were condition and log_2_ max(Ca_i_^2+^).

| Fig | variable | dend diam | F, p | T, P for $\text{Ca}_i^{2+}$ | T, P for condition |
| --- | --- | --- | --- | --- | --- |
| 3a | spatial extent | 1.2 | 364, $<1e-5$ | 17.9, $<1e-5$ | RyR2 no CaM: 21.1, $<1e-5$; no RyR2: -1.83, 0.069 |
| 3a | spatial extent | 2.4 | 122, $<1e-5$ | 16.0, $<1e-5$ | RyR2 no CaM: 7.4, $<1e-5$; no RyR2: -1.48, 0.142 |
| 3a | spatial extent | 6.0 | 148, $<1e-5$ | 17.7, $<1e-5$ | RyR2 no CaM: 9.6, $<1e-5$; no RyR2: -0.49, 0.867 |
| 3b | time decay | 1.2 | 63, $<1e-5$ | 11.2, $<1e-5$ | RyR2 no CaM: 8.7, $<1e-5$; no RyR2: 0.1, 0.934 |
| 3b | time decay | 2.4 | 48, $<1e-5$ | 5, $<1e-5$ | RyR2 no CaM: 9.0, $<1e-5$; no RyR2: -0.34, 0.734 |
| 3b | time decay | 6.0 | 66, $<1e-5$ | 11.9, $<1e-5$ | RyR2 no CaM: 5.8, $<1e-5$; no RyR2: -1.1, 0.267 |
| 5a | spatial extent | 1.2 | 434, $<1e-5$ | 23.8, $<1e-5$ | 13.7, $<1e-5$ |
| 5a | spatial extent | 2.4 | 344, $<1e-5$ | 22.2, $<1e-5$ | 11.1, $<1e-5$ |
| 5a | spatial extent | 6.0 | 368, <1e-5 | 25.6, <1e-5 | 5.7, <1e-5 |
| 5b | time decay | 1.2 | 68, <1e-5 | 7.6, <1e-5 | 7.6, <1e-5 |
| 5b | time decay | 2.4 | 78, <1e-5 | 7.6, <1e-5 | 8.9, <1e-5 |
| 5b | time decay | 6.0 | 185, <1e-5 | 12.4, <1e-5 | 13.0, <1e-5 |
| 6a | spatial extent | 1.2 | 470, <1e-5 | 31.5, <1e-5 | aging: 15.7, <1e-5; aging with ctrl RyR2: 2.4, 0.015 |
| 6b | time decay | 1.2 | 74, <1e-5 | 9.6, <1e-5 | aging: 10.1, <1e-5; ctrl+200% buffers: 3.1, 0.002; RyR2 no CaM: 4.3, <1e-5 |

**Fig 5.**
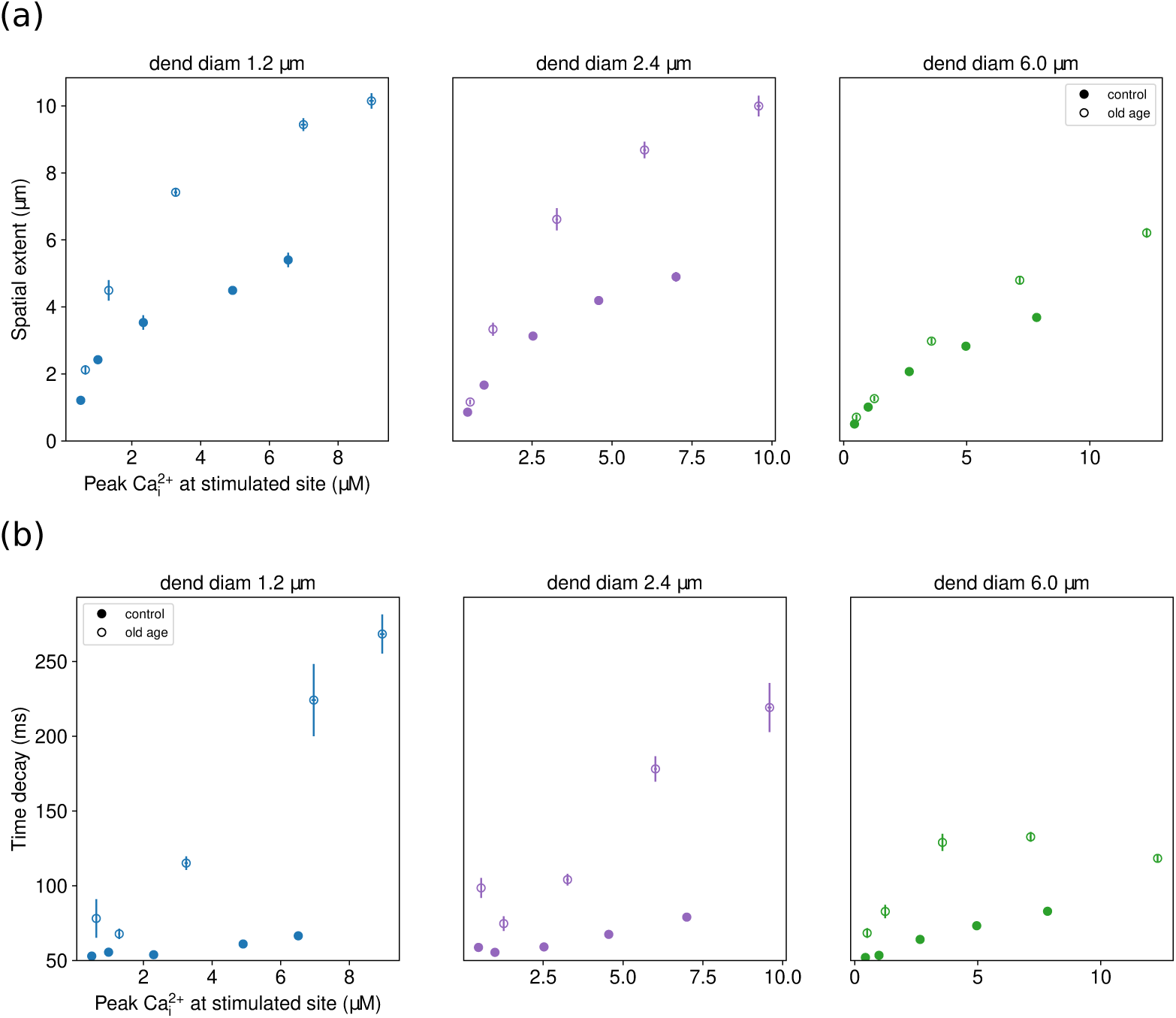
Old age prolongs calcium transients in space and time. The difference in spatial spread and temporal decay constant for the ctrl model (RyR2 inhibited by calmodulin) and old age model is statistically significant (*p <* 0.001 for every panel of (a) and (b)). (a) Spatial extent of calcium elevation in control conditions (filled circles) and the old-age model (empty circles). Spatial extent was the furthest location from stimulated site where calcium amplitude following stimulation exceeded 250% of resting calcium concentration. (b) Duration of calcium elevation in control conditions (filled circles) and the old-age model (empty circles). Time decay constant was obtained by fitting a single exponential decay function to calcium concentration after the stimulation at the stimulated section of the dendrite.

**Fig 6.**
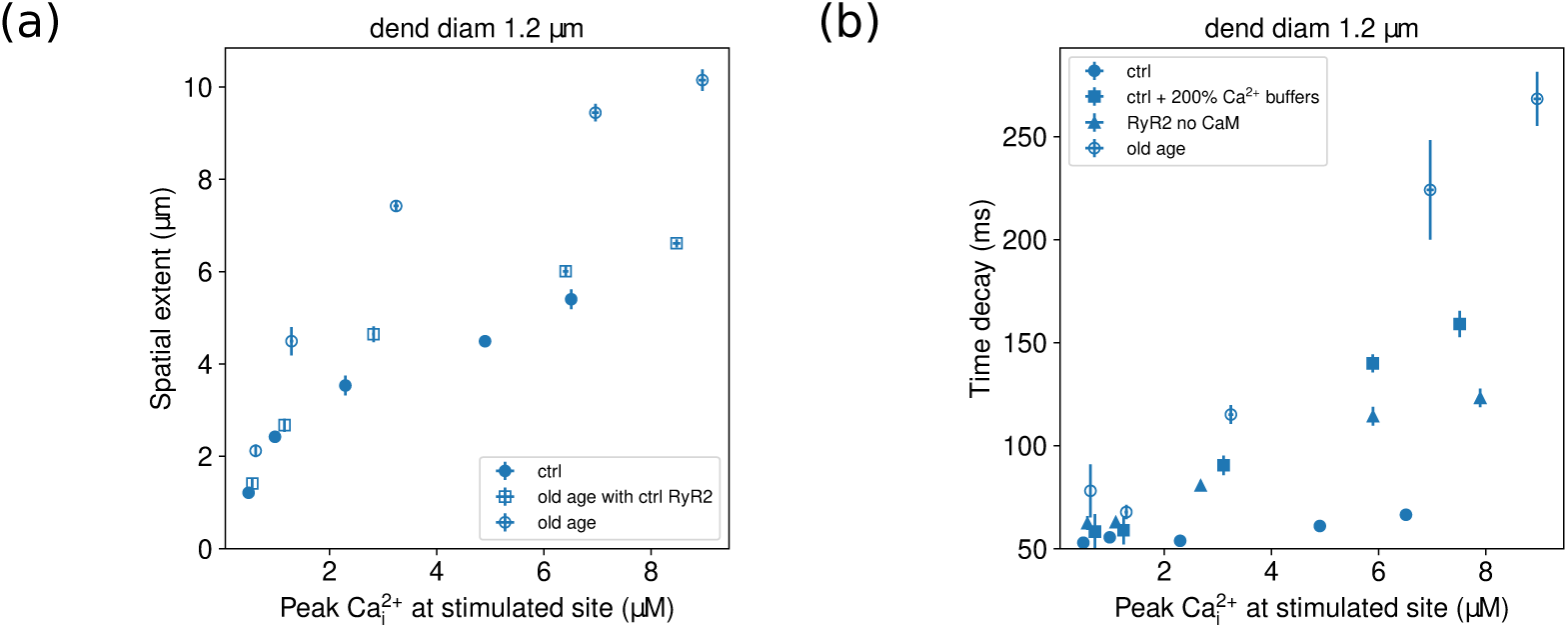
Comparison of spatial spread and temporal decay constant of calcium transients for models applying single changes from the old age model with the control model (solid circles) and the old age model (empty circles). Only results for the thin dendrite are shown. (a) Restoring RyR2 inhibition in the old age model (empty squares) lowered the spatial extent to levels similar to the ctrl model. Spatial extent was the furthest location from stimulated site where calcium amplitude following stimulation exceeded 250% of resting calcium concentration. (b) Either increasing Ca^2+^-buffering of the ctrl model (filled squares) or disinhibition of RyR2 (filled triangles) decreases temporal decay constant. Filled circles show the extent of temporal specificity in control conditions. Time decay constant was obtained by fitting a single exponential decay function to calcium concentration after the stimulation at the stimulated section of the dendrite.

The kymographs (Fig 7) show temporal dependence of the intracellular calcium concentration (averaged across the width) for every 0.5µm in the longitudinal direction. The mean and standard deviation of the intracellular calcium concentration (Fig 8) were calculated across the entire morphology for the whole simulation of 1000 s.

**Fig 7.**
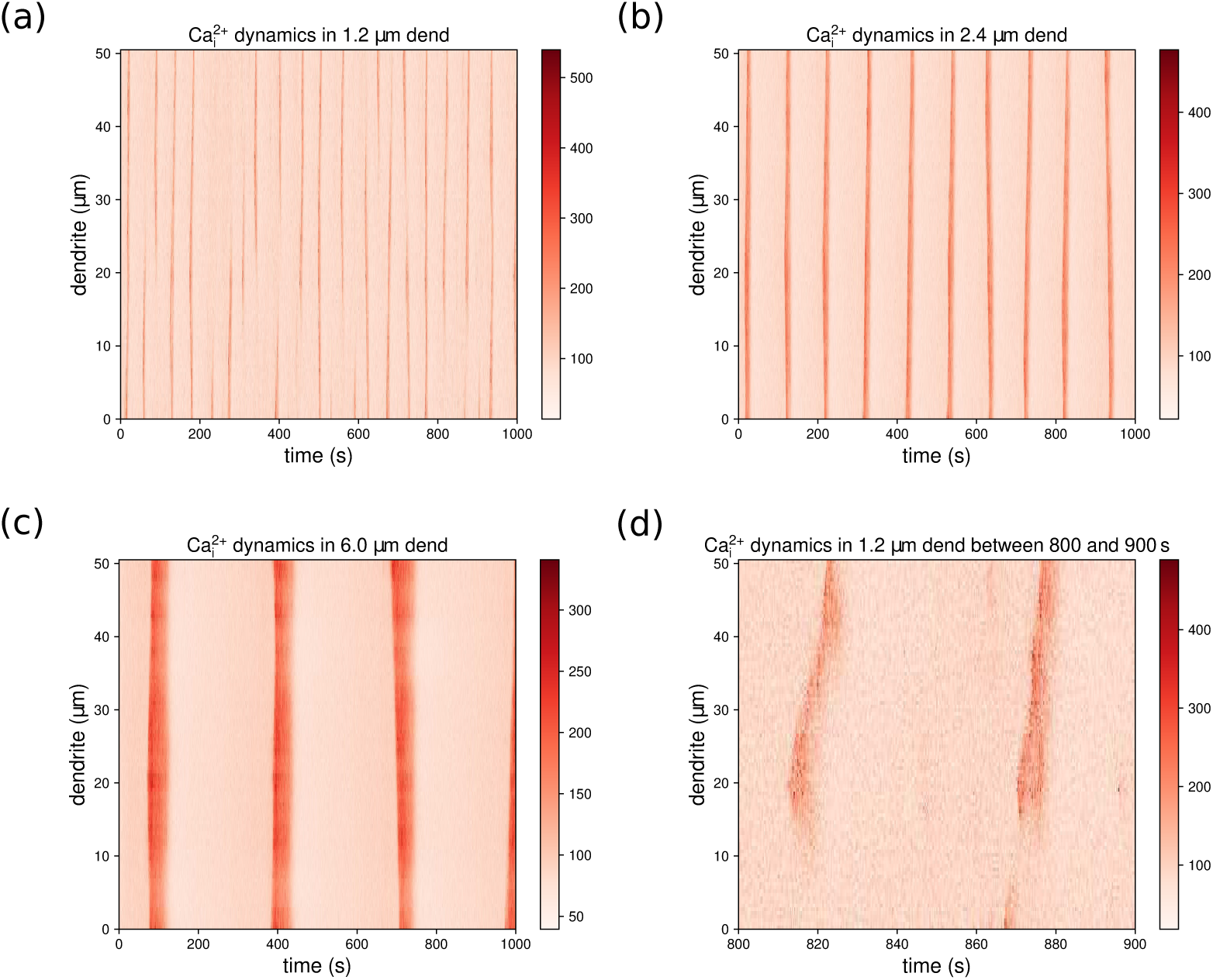
Spontaneous generation of calcium events in the AD model are seen in kymographs of typical resting intracellular calcium concentration (Ca_i_^2+^ in the AD model with (a) the 1.2 µm dendrite, (b) the 2.4 µm, and (c) the 6.0 µm dendrite. (d) Expansion of (a) showing that calcium waves sometimes begin in the middle of the dendrite.

**Fig 8.**
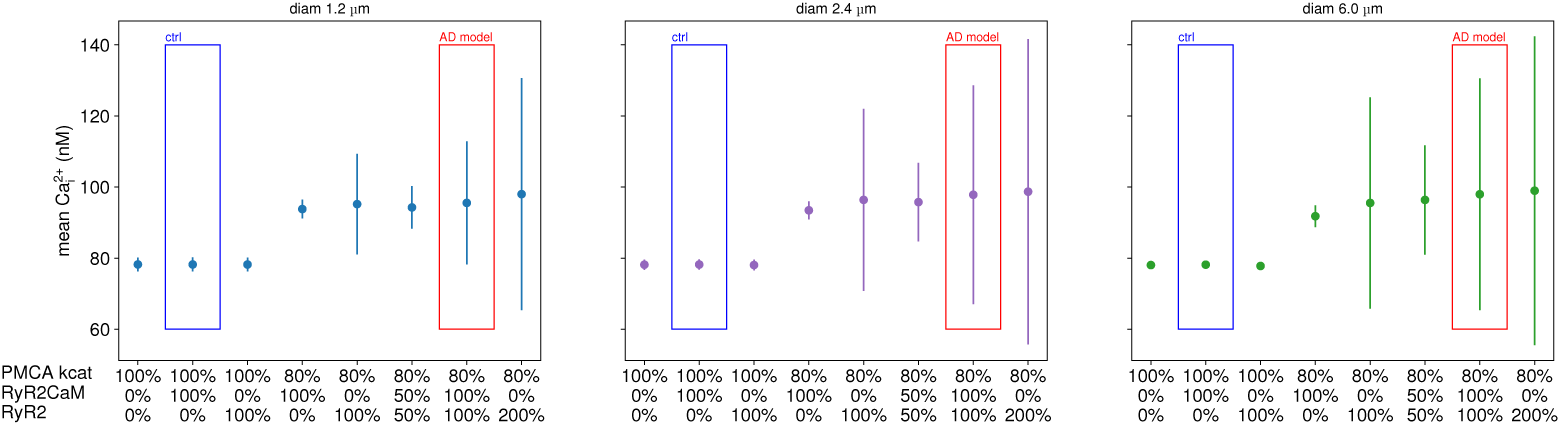
Lowering activity of PMCA pumps is necessary for an elevation in resting calcium concentration, while disinhibition of RyR2s increases the amplitude of calcium fluctuations. Each panel shows mean and standard deviation of intracellular calcium concentration Ca_i_^2+^ in the AD model (red box) and 7 models with varying PMCA activity and concentration of RyR2 inhibited by calmodulin (denoted as RyR2CaM) and disinhibted RyR2 (denoted as RyR2). The ctrl model is denoted with the blue box. RyR2 disinhibition alone did not change resting intracellular calcium concentration and can not explain the elevated resting intracellular calcium concentration observed in AD. Further increasing the concentration of disinhibited RyR2 (100% RyR2 concentration of the ctrl model) only increased resting intracellular calcium fluctuations. Mean and standard deviation of resting calcium concentration was calculated across all the time points and voxels in a 1000 s long simulation.

For Fig 9 we identified calcium events in the AD models as clusters, in space and time, of elevated intracellular calcium concentration, larger than 250% of the mean intracellular calcium concentration of the control model). Amplitude is the mean of maximum amplitude of the calcium events; peak frequency is the number of calcium events divided by the simulation duration (1000 s); peak duration is the average duration of the calcium events; and spatial spread is the average of spatial spread of the calcium events. Figure 9 shows mean values of the spatial spread and temporal decay contant with standard error as errors bars obtained in 4 simulations with different seeds.

**Fig 9.**
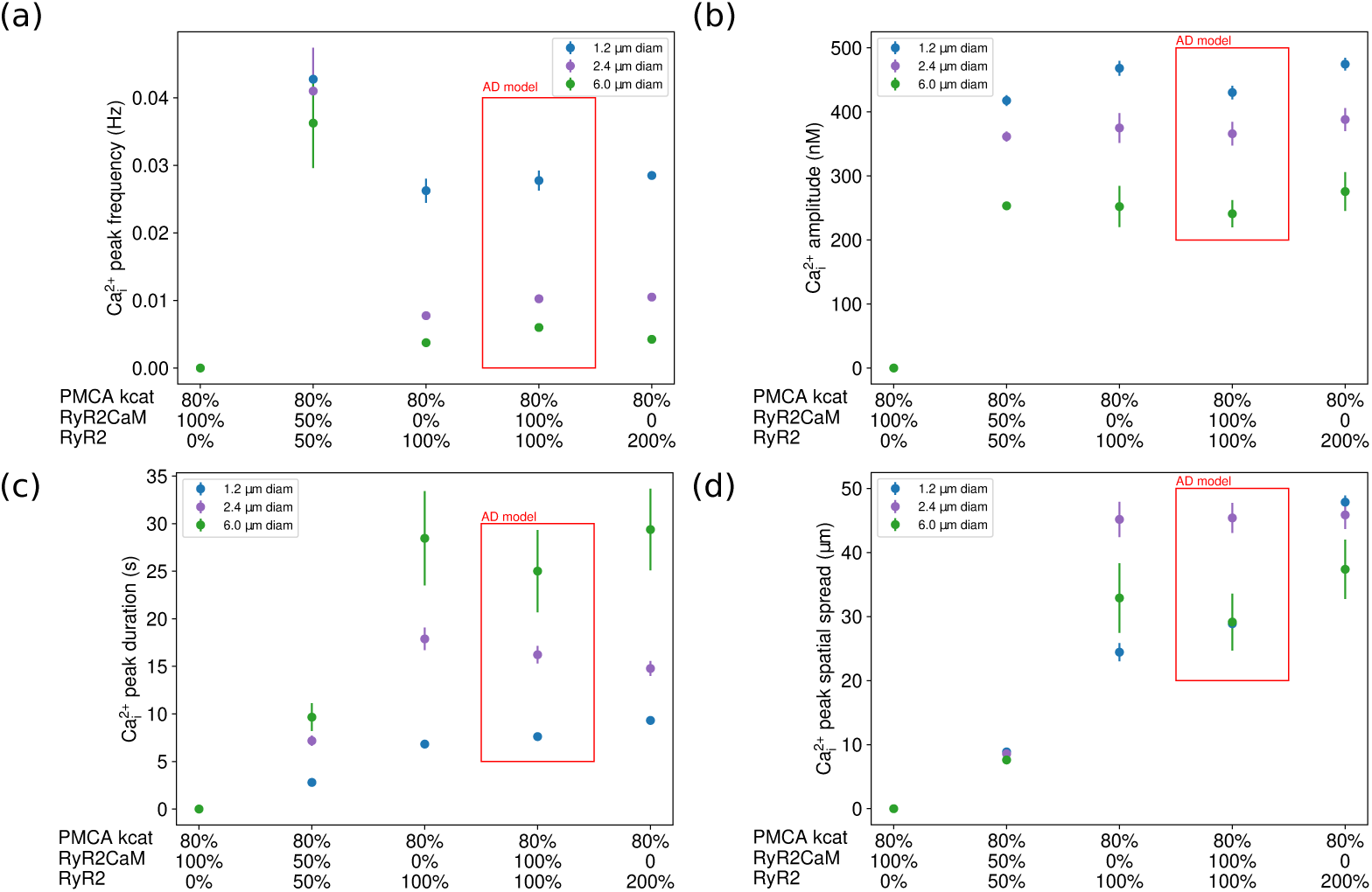
Properties of calcium events observed in the AD model (red box) and the 4 models with diminished PMCA activity. Calcium events were identified as a calcium elevation exceeding 250% of mean resting intracellular calcium concentration of the control model. (a) Frequency of calcium events was inversely proportional to the dendritic diameter. Frequency was calculated as the average number of calcium events divided by the duration of the simulation. (b) Amplitude of calcium events was inversely proportional to the dendritic diameter. Mean amplitude was calculated as the average maximum amplitude of the calcium events. (c) Peak duration of the calcium events was proportional to the dendritic diameter. Mean peak duration was calculated as the average duration of the calcium events. (d) There is not a consistent effect of diameter on the peak spatial spread. Mean spread was calculated as the average spatial spread of the calcium events.

To establish whether different model variants yield different spatial spread and temporal decay constant of calcium transients we performed linear regression analysis of spatial spread/temporal decay values versus log_2_ max(Ca_i_^2+^), which linearized the results. For all analyzed model variants (ctrl, no RyR2, RyR2 no CaM, old age, old age with RyR2CaM, ctrl with 200% Ca^2+^-buffers) linear regression fits were statistically significant with *p <* 0.001. To establish if model variants differ we performed ANCOVA, with model variant as co-variate, for every model panel in Figs 3, 5 and 6. For statistical analysis we used statsmodels Python package [86].

## Results

### RyR2s do not support intracellular calcium wave propagation

To test in-silico whether RyR2 alone can support intracellular calcium wave propagation caused by synaptic activation, we stimulated with calcium input models of a 51 µm long dendrite with three diameters: 1.2 µm (thin), 2.4 µm (medium) and 6.0 µm (thick). We modeled RyR2 constitutively bound to calmodulin (denoted as ctrl model), because the concentration of free calmodulin in the brain exceeds 10 µm [14, 15] and calmodulin affinity for the inhibitory binding site of RyR2 is in the range of 820 nM for apo-calmodulin [38] to 4 nM [65] for calcium-bound calmodulin. The binding of calmodulin to RyR2 is in fact very slow with dissociation rate *τ*1*_/_*_2_ reaching up to minutes [65] suggesting that most of RyR2s are bound to calmodulin in control conditions even during a calcium transient. To demonstrate the role of calmodulin inhibition, we also simulated the model without any RyR2.

We found that the ctrl model cannot support intracellular calcium wave propagation, and the absence of calmodulin-bound RyR2 (RyR2CaM) had no effect on the spatial extent of calcium transients in thin and medium dendrites (Fig 3a, Tab 11). Spatial spread of synaptically evoked calcium transients depended on the stimulation intensity, which was consistent with experimental observations [24] (Tab 11). The increase in spatial extent with stimulation intensity was lowest in thick dendrites, probably due to the larger impact of buffering [87]. Without RyR2 or with calmodulin inhibited RyR2, the spatial spread of calcium transients did not exceed 5 µm from the stimulation location, which is consistent with values reported in [87] (2.5 µm measured around the source as full width around half maximum). Stimulation intensity also prolonged calcium transients, again with no difference between excluding RyR2 and calmodulin inhibited RyR2.

### Calmodulin inhibition of RyR2 preserves spatial specificity of calcium transients

To investigate the role of calmodulin inhibition of RyR2 in shaping calcium dynamics, we substituted the model of RyR2CaM with a model of disinhibited RyR2 developed by [37]. We immediately observed an increase in calcium leak from the ER, which resulted in lowering of luminal calcium concentration by 10% in medium and thick dendrites. Resting intracellular calcium concentration levels stayed constant and excess intracellular calcium was extruded into the extracellular space by the PMCA and NCX.

Disinhibition of RyR2 increased the spatial spread (Fig 3) by a factor of 2 and greatly increase the temporal decay constant of the calcium elevation (Fig 3). Both effects were statistically significant (*p <* 0.001 for both spatial spread and temporal decay constant for every dendritic diameter, Tab 11). However, the increase in spatial spread did not allow for calcium wave propagation either for the synaptic-like (Fig 3) or short bAP-like stimulation (Fig 4).

For the disinhibited model both spatial and temporal specificity of calcium transients were lowest in thin dendrites and highest in thick dendrites.

### Calmodulin controls spatial and temporal specificity of calcium transients in old age

Old age is accompanied by larger [88] and longer [89] calcium transients with higher spatial spread [90], which lead to higher calcium accumulation in the cytosol [91], which is likely responsible for difficulties in learning and memory. Old age is also characterized by higher calmodulin oxidation [64], which in turn causes lower activity of PMCA pumps [66–68] and reduces calmodulin binding to RyR2 [16, 63, 92] resulting in RyR2 favoring activation by calcium [74]. RyR2 expression is also increased [71, 93]. Moreover, the buffering capacity of intracellular calcium buffers of CA1 pyramidal neurons is increased [70]. Finally in old age store-operated calcium entry [69] and the T-type VGCC [72] are down-regulated. To investigate how these changes interact to produce deficits in calcium dynamics we created an old age model implementing those changes.

First we simulated synaptic activation as a 40 ms calcium input into submembrane voxels for the model of old age (Fig 5). We saw that increased calmodulin oxidation (resulting in disinhibition of RyR2 and lower activity of PMCA pumps) in the old age model was responsible for decreased spatial and temporal specificity of calcium transients. The simulations showed that thick dendrites were the most robust to calmodulin oxidation (Fig 5a), although for all dendrite models, the increase of spatial spread in the old age compared to the ctrl model was statistically significant (Tab 11, *p <* 0.001). All dendrite models exhibited prolongation of the calcium elevation (Fig 5b) (Tab 11, *p <* 0.001) which is consistent with results from the previous section, although the duration of calcium transients was two times lower than for disinhibition of RyR2 alone.

Next we investigated which pathways downstream of calmodulin activation are responsible for the increased spatial extent (Fig 6a) and longer duration (Fig 6b) of calcium transients in old age. We investigated two separate aspects of the old age changes in calmodulin activity with the following parameter variations:

1. old age with ctrl RyR2: 80% of PMCA activity, 200% calcium buffering molecules concentration, 200% RyR2CaM,
2. ctrl with 200% of Ca^2+^-buffers: ctrl model with 200% calcium buffering molecules concentration,

and repeated in-silico experiments. Disinhibition of RyR2s was responsible for diminished spatial specificity of calcium transients. Fig 6a shows that the old age model with ctrl RyR2, i.e. inhibited by calmodulin (old age with ctrl RyR2, empty squares) exhibited similar spatial spread to the control model (filled circles) (Tab 11, *p* = 0.002).

Fig 6b indicates that both RyR2 disinhibition (RyR2 no CaM, filled triangles) and higher calcium buffering (ctrl+200% Ca^2+^-buffers, filled squares) each can increase duration times of calcium transients (Tab 11, *p* = 0.002), although this increase is lower than the one observed with the old age model. This is consistent with results shown in the previous section, which revealed that disinhibition of RyR2 significantly impacted temporal properties of calcium transients, and indicates that both these mechanisms contribute to diminished temporal specificity of calcium transients in the old age model.

### Disinhibition of RyR2 alone cannot account for increased resting intracellular calcium concentration in Alzheimer’s disease

Calcium dishomeostasis [76] is one of the hallmarks of Alzheimer’s disease [76]. One of the ways this dishomeostasis is manifested is an elevation in resting cytosolic calcium concentration, which is observed in pyramidal neurons of mice models of AD [77, 78]. This elevation may be caused by RyR2 recruitment [94], as indicated by an increase in CICR and higher RyR expression [79, 80]. Decreased inhibition of RyR2 by calmodulin can also cause increased RyR2 recruitment. Limiting RyR2 open time either by increasing its binding to calmodulin [1] or by a point mutation [7] alleviates cell loss and AD-like neuronal hyperexcitability, respectively. Alternatively, increased resting intracellular calcium concentration may be caused by a dysfunction of PMCA pumps [95], which are inhibited by tau proteins [96, 97] and *β*-amyloid oligomers [98], both of which are accumulated during AD progression. To investigate possible causes of elevated resting intracellular calcium concentration in AD we tested the impact of these implicated mechanisms in-silico using an early onset AD model, which comprised the ctrl model (Tab 2) with lower PMCA activity (Tab 8), increased RyR2 concentration (Tab 7) with half the RyR2 population inhibited by calmodulin and half disinhibited. It is important to note that the old age model and the AD model represent two different conditions. The old age model differs from the AD model in (a) the lack of SOCE and (b) higher buffering molecules, including neurogranin, and (c) unchanged calcium leak from the extracellular space, which represents resting calcium influx to the dendrite..

A typical kymograph of the AD model resting intracellular calcium shows spontaneous generation of spreading calcium elevations (Fig 7a) for the 1.2 µm model, and calcium elevations that traveled throughout the whole dendrite (calcium waves) for the 2.4 µm (Fig 7b) and 6.0 µm (Fig 7c) models. Amplitudes of these elevations exceeded 250% of the mean resting intracellular concentration in the control model (73 nM). These spontaneous events were not observed for the old age model, suggesting that in old age increased buffering helps preserve calcium homeostasis.

We varied the parameters to investigate which mechanisms are responsible for generation of the spontaneous calcium events:

1. 100% PMCA kcat 0% RyR2CaM 0% RyR2 – normal PMCA activity and no RyR2 receptors,
2. PMCA kcat 0% RyR2CaM 100% RyR2 – normal PMCA activity and all RyR2 receptors disinhibited,
3. 80% PMCA kcat 100% RyR2CaM 0% RyR2 – lower PMCA activity and all RyR2 receptors inhibited by calmodulin (AD without inhibited RyR2),
4. 80% PMCA kcat 0% RyR2CaM 100% RyR2 – lower PMCA activity and all RyR2 receptors disinhibited(AD without disinhibited RyR2),
5. 80% PMCA kcat 50% RyR2CaM 50% RyR2 – lower PMCA activity and half RyR2 receptor inhibited by calmodulin and half disinhibited (AD model with lower levels of both RyR2),
6. 80% PMCA kcat 0% RyR2CaM 200% RyR2 – lower PMCA activity, doubled RyR2 concentration and all RyR2 receptors disinhibited, and simulated them without any stimulation for 1000 s to evaluate their resting intracellular calcium concentration.

Fig 8 shows that RyR2 disinhibition alone did not increase resting intracellular calcium concentration. However, decreasing PMCA activity by 20% increased resting intracellular calcium concentration by 25% compared to the ctrl model. Interestingly, models with 80% of PMCA activity with increasing RyR2 concentration and increasing disinhibition of RyR2 did not show further increase of resting intracellular calcium concentration. Instead fluctuations of resting intracellular calcium concentration (represented by increased standard deviation of resting intracellular calcium concentration, shown in Fig 8) were observed. This shows that relieving calmodulin inhibition of RyR2s alone was not responsible for increased resting calcium concentration, but, instead, increased calcium fluctuations.

A closer look at cases summarized in Fig 8 revealed that emerging calcium events, some of which were calcium waves, were observed for models with diminished PMCA activity (denoted as 80% PMCA kcat), such as the model with 80% of PMCA activity and 50% of RyR2 disinhibition and 50% of RyR2CaM (this behavior was observed for the three dendrite models). Fig 9 summarizes the properties of the calcium events observed in all 5 models with diminished PMCA activity. The frequency of calcium events depended on the dendritic diameter (Fig 9a). With 50% of RyR2 disinhibited, calcium events were more frequent (Fig 9a), but also more localized (Fig 9a) and Fig 9c). The amplitude of calcium events depended only on the dendritic diameter (Fig 9b). Calcium events in thick dendrites had the longest duration (Fig 9b), but the largest spatial spread of calcium waves was observed in medium dendrites (Fig 9d).

### Store operated calcium entry does not affect CICR spatial specificity in control conditions

Finally we investigated the role of store replenishment in spatial specificity of CICR. The endoplasmic reticulum membrane contains STIM proteins, which serve as ER calcium sensors. When ER calcium is depleted, STIM molecules bind and activate Orai1 channels located in the plasma-membrane. Calcium flows into the cell through those channels and gets transported by SERCA pumps into the ER. Fig 10 shows that store operated calcium entry (SOCE) does not increase the spatial extent of calcium transients due to the much slower time scale of SOCE activation than that of calcium dynamics. Although, experimental evidence [99] shows that STIM-Orai1 signaling in the spine is necessary for maintaining long-term structural plasticity, in this work we utilized stimulation resembling single solitary synaptic activation, which barely affects luminal calcium levels.

**Fig 10.**
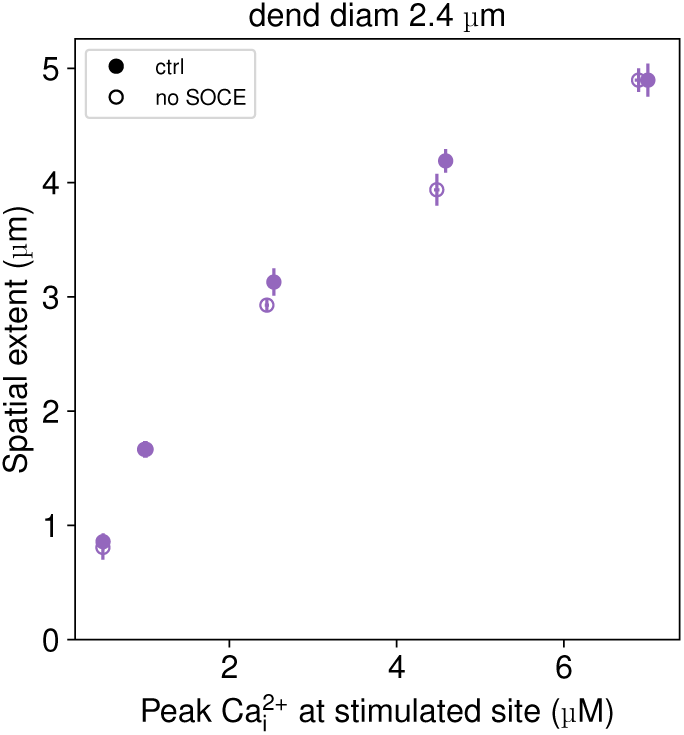
Store operated calcium entry (SOCE) did not influence spatial specificity of calcium elevations. Spatial extent was the furthest location from stimulated site where calcium amplitude following stimulation exceeded 250% of resting intracellular calcium concentration.

### Robustness of the model

We analyzed the change in spatial spread of calcium elevations in response to small changes in PMCA activity and concentration, and the luminal calcium concentration (shown in Fig 11). As expected due to the large impact of calcium buffering, thick dendrites were the most robust to parameter variation. Small increases in ER calcium concentration did not affect the spatial extent of calcium elevations for all three model dendrites. PMCA concentration and activity changes impacted thin and medium dendrites the most.

**Fig 11.**
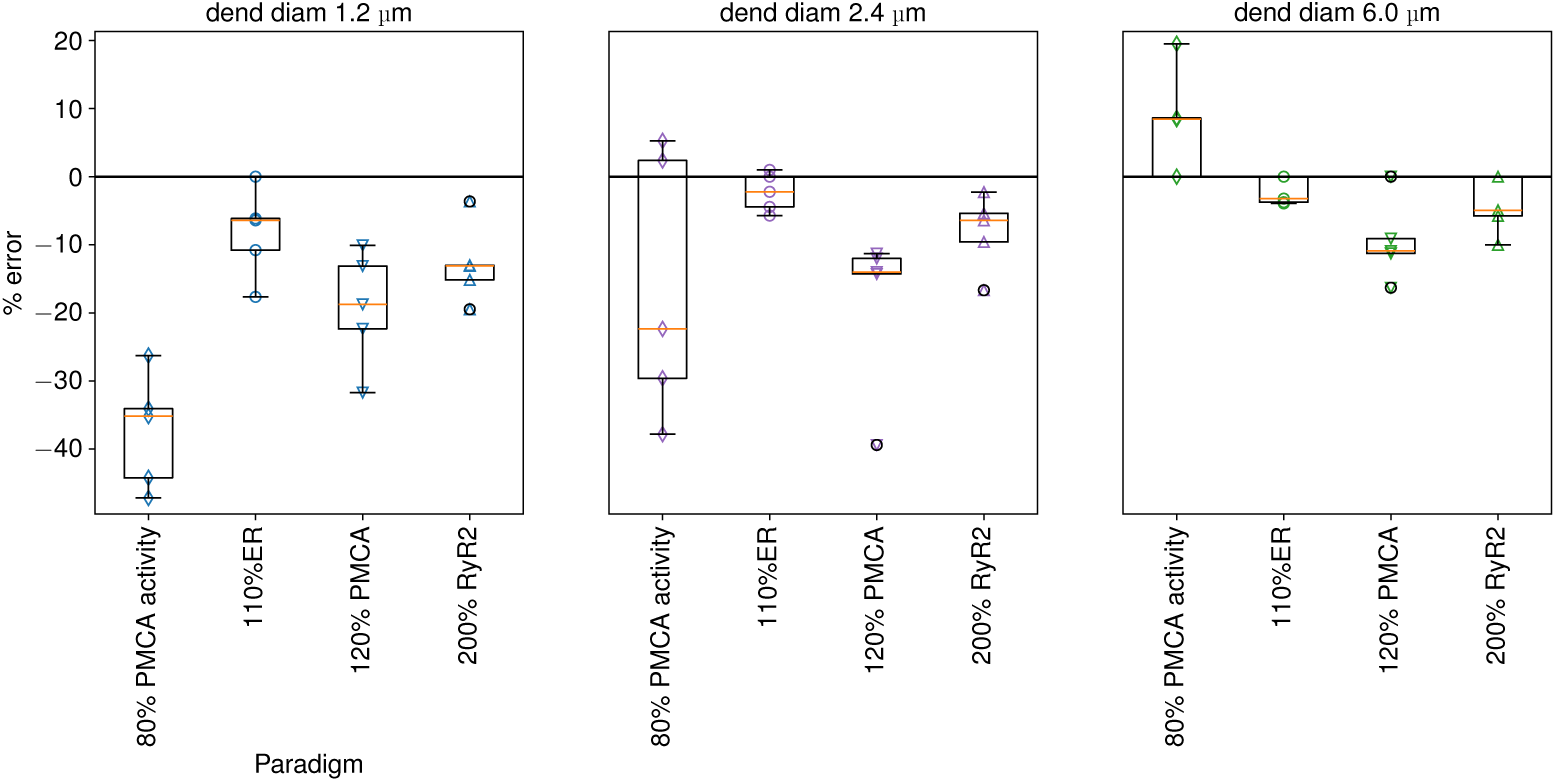
Thick dendrites are most robust to parameter variation. Panels illustrate the change in spatial extent for the range of calcium influx values. Change in PMCA activity produced the highest variation in spatial extent, especially for thin and medium dendrites. Horizontal lines represent mean and standard deviation of percentage error for each paradigm. Open symbols represent individual values of percentage error.

## Discussion

In this work we investigated how calcium homeostasis is controlled by calmodulin in control conditions, aging and Alzheimer’s disease. We concentrated on calcium-induced calcium release, in which calcium flows out of the ER into the cytosol through RyR2, and calcium extrusion into the extracellular space by PMCA. We showed that RyR2s alone are unlikely to support intracellular calcium wave propagation both in control conditions (when they are constitutively bound to and in consequence inhibited by calmodulin) and in old age (when both RyR2s and calmodulin become oxidized, which in turn disinhibits some of the RyR2s). Calmodulin inhibition of RyR2s played a crucial role in controlling spatial and temporal specificity of calcium transients. Finally, we investigated possible sources of increased resting intracellular calcium concentration in Alzheimer’s disease, showing that disinhibition of RyR2s could not account for this phenomenon, but, instead, diminished activity of PMCA increased intracellular resting calcium.

The influence of calmodulin and RyR2 on calcium dynamics is critical because learning, memory and many other processes in the brain are controlled by calcium dynamics. Dendritic calcium plays a crucial role in behavioral time scale plasticity, a very powerful form of synaptic plasticity requiring a prolonged dendritic calcium elevation, which underlies tuning of spatial features in dendrites and is capable of producing place cells [100]. Interestingly, blocking mitochondrial calcium buffering, and hence increasing calcium release from the ER, led to higher stability of place cells [101]. Dendritic calcium is strongly correlated with spine calcium [84] with RyR3s (activated by calmodulin) on the spine endoplasmic reticulum (spine apparatus) transducing spine calcium signals to the dendrite [102, 103]. Calmodulin acting on RyRs differentially regulates the calcium transients in the dendritic branch: leading to an amplification of calcium transients from individual spines and restricting their spread in the dendrite leading to larger spatial and temporal specificity of calcium signaling in the dendrite. Thus, RyRs can influence learning and memory through their contribution to dendritic and spine calcium transients.

Results showing the lack of intracellular calcium wave induction in the control model are in agreement with experimental evidence, which showed that IP_3_ production is necessary [24–27], and in conflict with other modeling work [23], which did not incorporate calmodulin inhibiting RyR2. More recent work in hippocampal cultures shows that RyR2 signaling contributes to nuclear calcium release [21], although this process required depolarization of membrane potential [19, 20], which was not included in the model. Moreover, contrary to [23] in Fig 3 we showed that, even if RyR2s were disinhibited by calmodulin, calcium inputs corresponding to excitatory postsynaptic potentials (EPSP) and bAPs (Fig 4) could not elicit intracellular calcium waves. The lack of calcium waves may be advantageous since limiting intracellular calcium release supports the establishment of dendritic feature selectivity [101].

Although, we used similar RyR2 densities (4 µm^−2^) in our model, there were many methodological differences from [23]: instead of using a deterministic model we used a stochastic approach for simulation of reaction-diffusion systems, which is better suited to dynamical systems with low copy numbers [104] and in such systems deterministic models might yield different predictions than stochastic models [105]. One of the differences is that deterministic rate laws depend on concentrations whereas stochastic rate laws depend on the number of possible combinations of reacting species. With high copy numbers stochastic and deterministic models will yield similar results, whereas with low copy numbers the reactions might differ substantially, e.g. contributing to higher noise sensitivity of deterministic models [105]. The stochastic approach we used is a spatial extension of Gillespie’s tau leap algorithm, which dynamically selects between simulating an exact event or a leap event both for reactions and diffusion between voxels [85]. This algorithm provides the computational efficiency needed for simulating this mathematically stiff system. Finally, we used a stochastic model of RyR2 activation [37] tuned to match mean open and close time of the RyR2 [39] instead of using the Keizer-Levine model of RyR2 activation [106], which exhibits very long RyR2 open times. It is possible that the induction of intracellular calcium waves in [23] resulted from using the Keizer-Levine RyR2 model. In fact, when we performed stochastic simulations with the model of resting dynamics, we observed spontaneous, oscillatory generation of intracellular calcium waves in thin dendrite. We did not compare predictions of our model with [28] and [29], because both models implemented IP_3_-dependent calcium wave generation.

We modeled calmodulin constitutively bound to RyR2s in control conditions. Calmodulin concentration in the hippocampus is high (20 µM) [14]) and calmodulin affinity for the inhibitory binding site of RyR2 in the range of 820 nM for apo-calmodulin [38] to 4 nM [65], with dissociation rate *τ*1*_/_*_2_ reaching up to minutes [65]. This suggests that most of RyR2s are bound to calmodulin in control conditions even during a calcium transient. In fact, we did a limited number of simulations with dynamic binding of calmodulin to RyR2 during a transient calcium elevation and observed that new calmodulin did not bind to the RyR2 subunit due to the slow reaction rates. In oxidized conditions and with cytosolic calcium levels (100 nM), calmodulin affinity for RyR2 drops twenty-fold [65] with 50% RyR2s bound to calmodulin. Therefore we simulated calmodulin constitutively bound to RyR2 in control conditions and 50% of RyR2 constitutively bound to calmodulin in the model of aging. This resulted in total RyR2 population favoring activation by calcium for the model of aging consistent with experimental data [74].

The endoplasmic reticulum in dendrites forms a dense network of interconnected cysternae with an occasional single large cysternae close to dendritic membrane with mitochondria in the middle of the dendrite [35, 107]. Modeling such morphology with fidelity would require a particle-based approach, which is computationally prohibitive in the scale of model used here. Instead, in our mesoscopic model of calcium dynamics in a 51 µm long dendrite, the ER was treated as a separate compartment that is uniformly distributed in the dendrite, occupying 25 % of its volume. Tortuosity of the ER was taken into account by reducing the diffusion rate of calcium molecules in the ER, compared to diffusion rate of the cytosolic calcium molecules. The randomly placed ryanodine receptors allow for a release of ER calcium molecules into the cytosol, with flow of calcium down the concentration gradient through open RyR receptors.

We showed that both spatial [90] and temporal specificity of calcium transients decreased due to the higher oxidation of calmodulin in aging, which likely contributes to deficits in learning and memory accompanying old age. In control conditions the extent of calcium elevation in thin dendrites reached 5 µm, which was similar to the average inter-spine distance after contextual fear conditioning task observed in apical dendrites of layer V motor neurons [108], indicating that this distance might indeed be governed by calcium release from the ER. When calmodulin inhibition of RyR2 was relieved (e.g. by higher oxidation of calmodulin), the spatial extent of synaptically evoked calcium transients was increased by roughly a factor of 2 and duration was increased for thin and medium dendrites. The decrease in spatial specificity was caused by the oxidized calmodulin not inhibiting RyR2, but the decrease in temporal specificity was caused both by RyR2 disinhibition and an increase in cytosolic calcium buffering. Interestingly, increased calcium cytosolic buffering acted like a double edged sword: it counteracted some of the calcium dishomeostasis accompanying old age and prevented spontaneous generation of calcium events and waves keeping the resting intracellular concentration unchanged [109], but also decreased temporal specificity of calcium transients.

Increasing oxidation levels leads to both lower PMCA activity [110] and RyR2 disinhibition [6, 16]. In our model of early onset AD, with unchanged resting calcium influx to the dendrite lower PMCA activity alone resulted in the elevation of intracellular calcium concentration. Coupling the lower PMCA activity with disinhibited RyR2 caused the emergence of spontaneous calcium events, with frequency dependent on dendritic diameter and disinhibited RyR2 concentration. In rat hippocampal CA1 pyramidal neurons [111–113] and CD1 mice hippocampal interneurons [114] spontaneous fast calcium events (calcium sparks) depended on RyR activation, resting intracellular calcium concentration and L-type VGCCs. The events observed in the model did not resemble that experimental phenomenon, instead their frequency was much lower (less than 0.02 event/sec compared to 1 event/sec [111, 112])) and their duration much longer (tens of second compared to 400 ms [112]). These results suggest that calcium events were driven by the degree of fluctuations caused by the increased activity of disinhibited RyR2s, which had a higher affinity for calcium ions, and elevated resting intracellular calcium concentration caused by lower activity of PMCA pumps.

A crucial question is whether disinhibition of RyR2s can be responsible for the elevated resting intracellular calcium concentration observed in Alzheimer’s disease [1, 7]. It is important to note that the old age model and the AD model represent two different conditions. The old age model differs from the AD model in (a) the lack of SOCE, (b) higher buffering molecules, including neurogranin, and (c) unchanged calcium leak from the extracellular space, which represents resting calcium influx to the dendrite. We showed that disinhibition of RyR2s, which was characterized by higher opening probability with longer open times [38, 39], caused higher calcium leak from the ER. This did not elevate resting intracellular calcium, because excess calcium was extruded outside the cell by the calcium extrusion pumps. We also investigated an alternative hypothesis of dysfunction of PMCA pumps [95] by lowering their activity, which resulted in an elevation of resting intracellular calcium concentration by 25% compared to the ctrl model. This increase was lower than the observed doubling of resting intracellular calcium concentration [49, 77, 78]. One limitation of the model is that it did not include *β*-amyloid. Calcium promotes *β*-amyloid oligomer formation [115], which can promote oxidative stress in neurons [116, 117]. Incubation of primary neuronal cultures with *β*-amyloid peptides leads to a significant increase in resting intracellular calcium and cell apoptosis [118, 119], which was prevented by blocking RyRs. These results suggested that lowering of PMCA activity in the vicinity of *β*-amyloid plaques [98] might be responsible for elevated calcium [78]. It is possible that in early asymptomatic stages of AD soluble *β*-amyloid oligomers promote calcium dishomeostasis via both inhibition of PMCA pumps and disinhibition of RyR2s, which in turn promotes *β*-amyloid plaque formation. Future models can evaluate whether a positive feedback loop between *β*-amyloid oligomer formation and calcium elevation can completely explain the experimentally observed doubling of resting calcium concentration.

## Conclusion

Calmodulin, the jack of many trades, controls calcium dynamics and homeostasis not only by acting as a fast calcium buffer: spatial and temporal specificity of calcium transients in dendrites of CA1 pyramidal neurons is regulated by calmodulin inhibition of RyR2 and basal intracellular calcium concentration by calmodulin activation of PMCA pumps. It is rather difficult to pinpoint one molecule responsible for calcium dishomeostasis. Instead, it is an ensemble of factors and molecular species interacting with each other with an emerging role for calmodulin, which regulates both calcium extrusion and calcium-induced calcium release.

## Supporting information

**S1 Fig 1. Spatial spread of calcium transients for a bAP-like stimulus.** A short calcium input (3 ms) was applied directly to the cytosol at one side of the dendrite as in [23] (also shown in Fig 2). Disinhibition of RyR2 by calmodulin (empty circles) increases spatial extent, but does not allow for calcium wave propagation. Spatial extent was the furthest location from the stimulated site where calcium amplitude following stimulation exceeded 250 % of resting calcium concentration. The horizontal error bars do not appear because they are smaller than the data points on the plots.

**S1 Fig 2. Comparison of spatial spread of calcium transients for models applying single changes from the old age model with the control model (solid circles) and the old age model (empty circles).** Restoring RyR2 inhibition in the old age model (empty squares) lowered the spatial extent to levels similar to the ctrl model. Spatial extent was the furthest location from stimulated site where calcium amplitude following stimulation exceeded 250 % of resting calcium concentration.

**S1 Fig 3. Comparison temporal decay of calcium transients for models applying single changes from the old age model with the control model (solid circles) and the old age model (empty circles).** Either increasing Ca^2+^-buffering of the ctrl model (filled squares) or disinhibition of RyR2 (filled triangles) increases temporal decay. Filled circles show the extent of temporal specificity in control conditions. Time decay constant was obtained by fitting a single exponential decay function to calcium concentration after the stimulation at the stimulated section of the dendrite.

## Acknowledgments

The authors would like to thank prof. Daniel K Wójcik for his continuous support.

## Notes

https://github.com/asiaszmek/stochastic_ER.git

### Competing Interest Statement

The authors have declared no competing interest.

### Summary of Updates

This is a revised version of the manuscript submitted to the journal.

