## Supporting material S1 for "Calmodulin controls spatial and temporal specificity of calcium-induced calcium release"

Supporting information

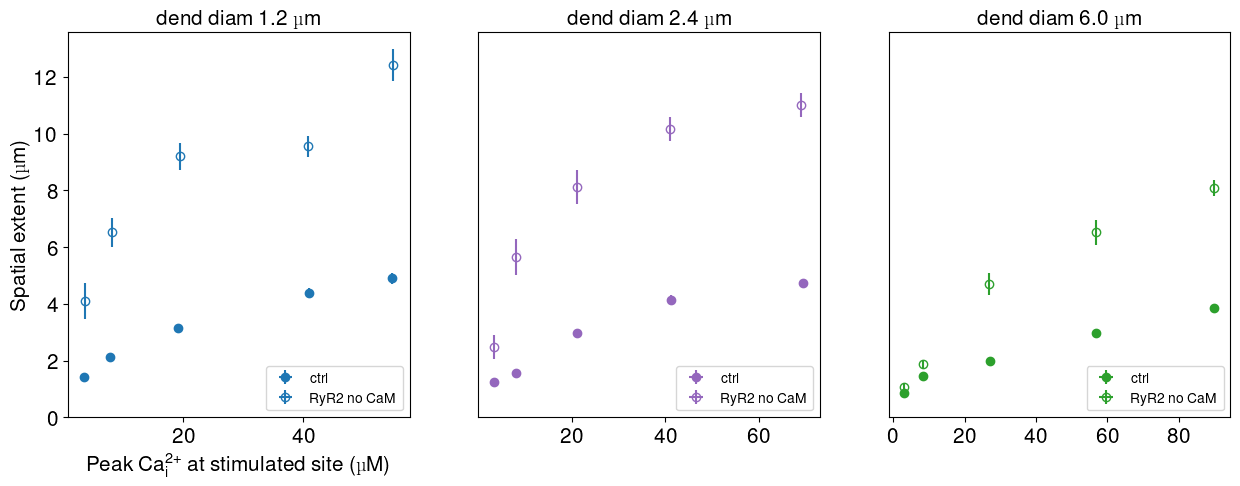
S1 Fig 1. Spatial spread of calcium transients for a bAP-like stimulus. A short calcium input (3 ms) was applied directly to the cytosol at one side of the dendrite as in [21] (also shown in Fig 2). Disinhibition of RyR2 by calmodulin (empty circles) increases spatial extent, but does not allow for

calcium wave propagation. Spatial extent was the furthest location from the stimulated site where calcium amplitude following stimulation exceeded 250% of resting calcium concentration. The horizontal error bars do not appear because they are smaller than the data points on the plots.

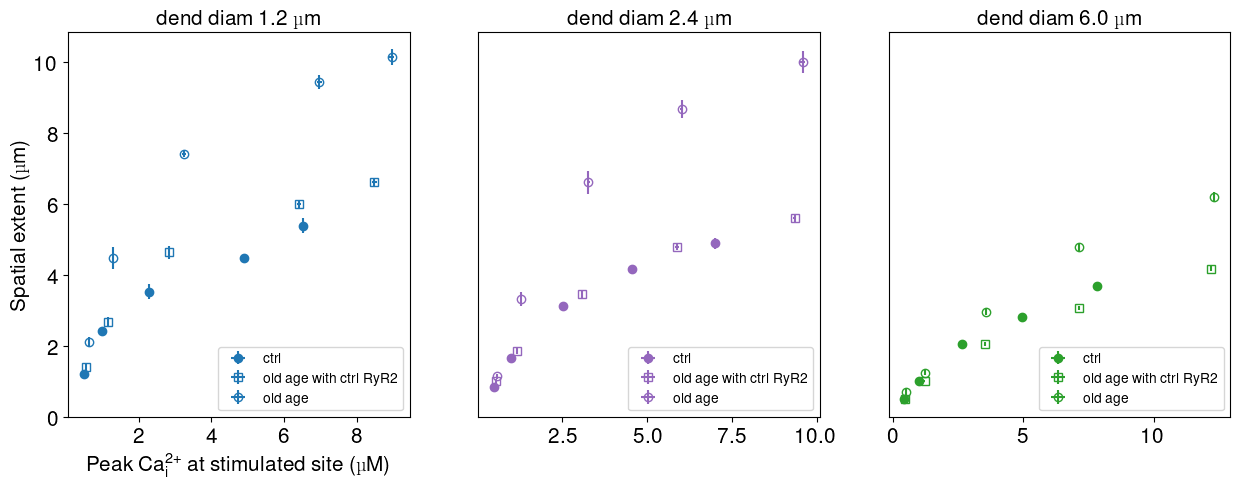
S1 Fig 2. Comparison of spatial spread of calcium transients for models applying single changes from the old age model with the control model (solid circles) and the old age model (empty circles). Restoring RyR2 inhibition in the old age model (empty squares) lowered the spatial extent to levels similar to the ctrl model. Spatial extent was the furthest location from stimulated site where calcium amplitude following stimulation exceeded 250% of resting calcium concentration.

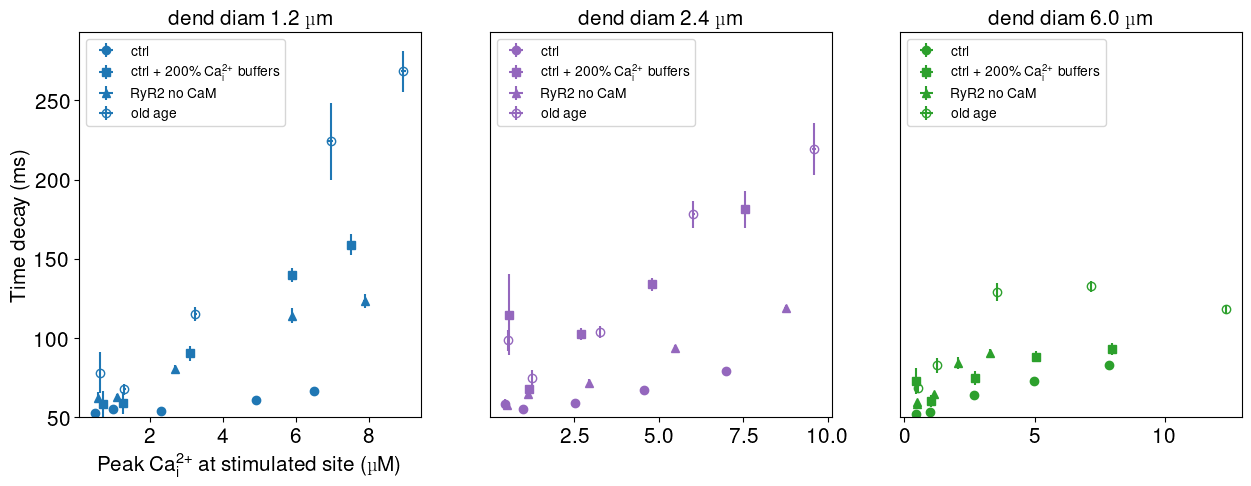
S1 Fig 3. Comparison temporal decay of calcium transients for models applying single changes from the old age model with the control model (solid circles) and the old age model (empty circles). Either increasing Ca2+ -buffering of the ctrl model (filled squares) or disinhibition of RyR2 (filled triangles) increases temporal decay. Filled circles show the extent of temporal specificity in control conditions. Time decay constant was obtained by fitting a single exponential decay function to calcium concentration after the stimulation at the stimulated section of the dendrite.
